# The curses of performing differential expression analysis using single-cell data

**DOI:** 10.1101/2024.05.28.596315

**Authors:** Chih-Hsuan Wu, Xiang Zhou, Mengjie Chen

**Affiliations:** Department of Statistics, University of Chicago, Chicago, USA; Department of Biostatistics, University of Michigan, Ann Arbor, USA; Department of Human Genetics and Department of Medicine, University of Chicago, Chicago, USA

**Author notes:** Corresponding authors: Correspondence to Mengjie Chen,.

## Abstract

Differential expression analysis is pivotal in single-cell transcriptomics for unraveling cell-type– specific responses to stimuli. While numerous methods are available to identify differentially expressed genes in single-cell data, recent evaluations of both single-cell–specific methods and methods adapted from bulk studies have revealed significant shortcomings in performance. In this paper, we dissect the four major challenges in single-cell DE analysis: normalization, excessive zeros, donor effects, and cumulative biases. These “curses” underscore the limitations and conceptual pitfalls in existing workflows. In response, we introduce a novel paradigm addressing several of these issues.

---

Differential expression (DE) analysis in single-cell transcriptomics provides essential insights into cell-type–specific responses to internal and external stimuli^1–4^. While many methods are available to identify differentially expressed genes from single-cell transcriptomics, recent studies raise important concerns about the performance of state-of-the-art methods, including both methods tailored to single cell data and techniques that work well in bulk^5–7^. As population-level single-cell studies rapidly become more feasible, powerful and accurate analytical methods will be essential for obtaining meaningful results. In this context, we discuss the four “curses” that currently plague the differential expression analysis of single-cell data: normalization, zeros, donor effects, and cumulative biases, highlighting the various limitations and conceptual flaws in the current workflows. We demonstrate these limitations using real data from 10X single-cell RNA-seq (sRNA-seq) data from post-menopausal fallopian tubes^8^. Finally, we present a new paradigm that offers a potential solution to some of these issues and illustrate its performance using two case studies.

## The curse of normalization

The term ‘normalization’ has been used to denote multiple distinct approaches in genomics^9, 10^. For example, it can refer to the process of correcting PCR amplification biases introduced during sequencing library preparation (library size normalization)^11–13^, the process of harmonizing data across different experimental batches (batch normalization)^14–18^, or to the process of transforming the data to adhere to a normal distribution (data distribution normalization)^19^. All three have been introduced to handle both bulk and single cell RNA-seq data, aiming to minimize unwanted technical variations. Choosing appropriate normalization techniques for DE analysis of scRNA-seq data is clearly important to maintain the integrity of the data, but the field has yet to establish a definitive gold standard outlining the circumstances for which different normalizations should be performed.

Library size normalization is critical in bulk RNA-seq analysis, as it is impossible to track the absolute abundance of RNA molecules in typical bulk RNA-seq protocols due to an unknown fold of amplification introduced by PCR during library construction. Normalization, in this instance, focuses on estimating and subsequently correcting for a sample-specific size factor. This process allows bulk RNA-seq to estimate relative RNA abundances. Post-normalization, samples are calibrated against a common reference, resulting in most genes displaying similar expression levels across samples. When performing differential expression analysis with bulk RNA-seq data, genes are classified as either up-regulated or down-regulated, based on the assumption that the majority remain unchanged across groups. While this size-factor based normalization technique is suitable for bulk RNA-seq, it does not translate effectively to scRNA-seq. Protocols in scRNA-seq, such as the 10X, employ unique molecular identifiers (UMIs) which discern between genuine RNA molecules and those generated via PCR. This enables the absolute quantification of RNA levels. Unfortunately, size-factor–based normalization methods, like counts per million reads mapped (CPM) convert data into relative abundances erasing useful data provided by the UMIs. Furthermore, CPM-normalized data does not account for competition among genes for cellular resources because the uniform number of molecules found in CPM-normalized data does not accurately represent true expression levels, which ultimately leads to suboptimal DE analysis results.

In batch effect normalization, dimension reduction methods pinpoint genes with consistent expression patterns across various batches; these genes act as anchors, guiding the alignment and integration of data^20^. However, in scRNA-seq analysis, only highly expressed or highly variable genes are retained for estimating batch effects and subsequent integration. As a result, gene numbers in integrated scRNA-seq datasets are noticeably reduced compared to the raw UMI data.

For data distribution normalization, the field offers both straightforward (e.g., log-transformation) and advanced strategies (e.g., variance stabilizing transformation, or VST). A notable implementation for scRNA-seq of VST is sctransform^21^, which employs a regularized negative binomial regression model, preserving the Pearson residuals for future analytical steps, including DE analysis^22^. However, if the underlying data distribution deviates significantly from the assumed model, the application of VST may introduce bias into the analysis.

To demonstrate the effects of various normalization methods on single-cell data, we compared the raw UMI counts of 10x scRNA-seq data obtained from post-menopausal fallopian tubes (see Methods) with data normalized using one of three methods: 1) CPM; 2) integrated log-normalized counts after removing batch effects using the Seurat CCA model^23^; and 3) VST data using sctransform^21^. As a result, we see the total UMI counts revealed substantial variations in library sizes across different cell types; notably macrophages (MP) and secretory epithelial (SE) cells exhibited significantly higher RNA content than other cell types (Fig. 1a). Furthermore, SE cells exhibited larger mean library sizes than mast (MA) cells across all donors. These findings align with the understanding that the main active cell types in post-menopausal fallopian tubes are MP and SE cells, with other cell types remaining dormant post-menopause. However, in the integrated data, the disparities in library size distribution were mitigated, even within cell types (Fig. 1a). While integration reduced differences across donors, it came at the cost of diminishing variation across cell types. It is worth mentioning that CPM normalization equalizes library sizes across all cell types; such normalizations may potentially obscure differences between cell types that are vital for understanding their unique biological functions.

**Figure 1.**
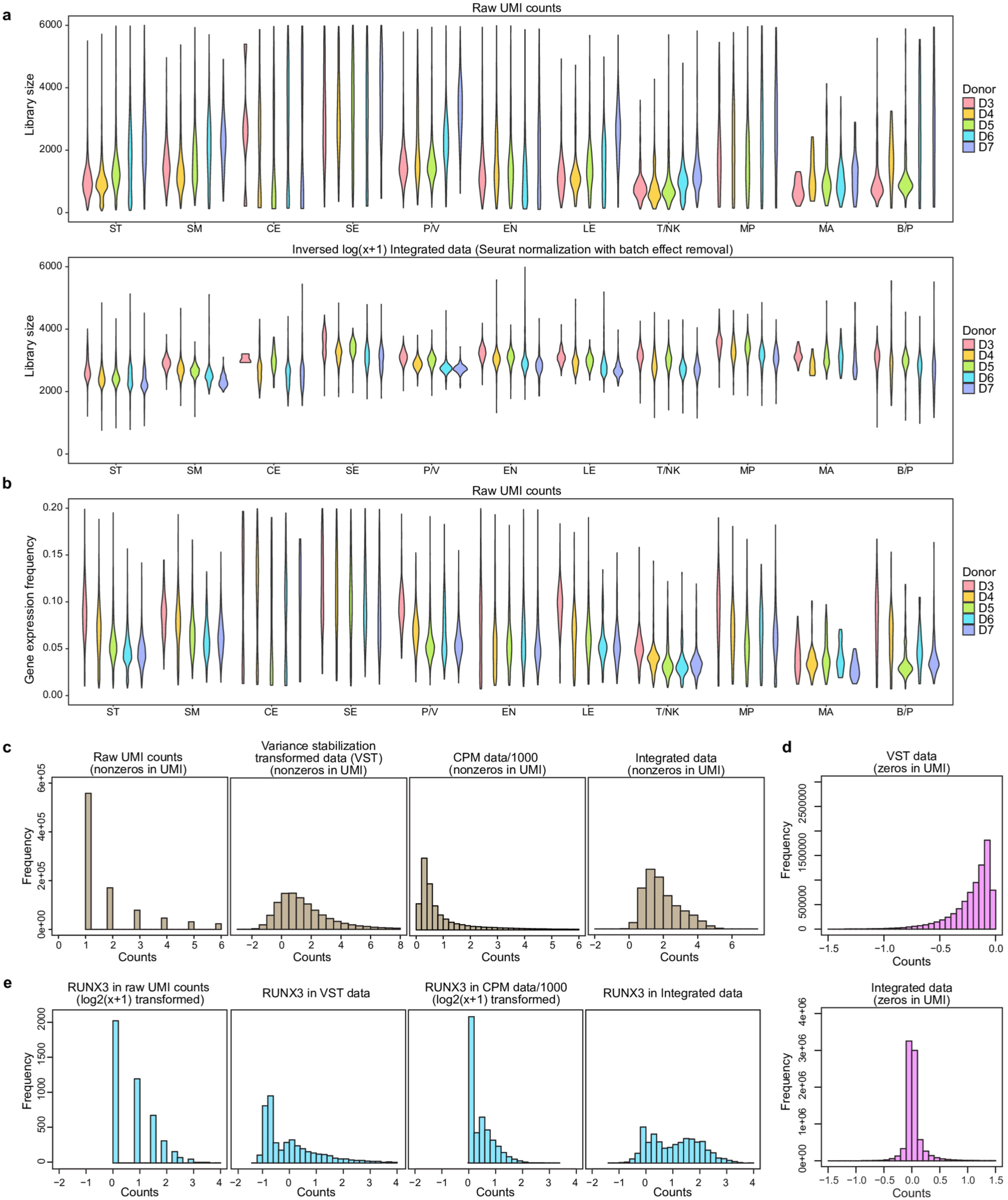
Effects of normalization on library size, zero frequency, and gene count distributions. a. Violin plots display library sizes based on raw UMI counts (top) and after data integration (bottom), categorized by cell types and donors. b. Violin plot illustrating the frequency of gene expression (non-zero counts) in raw UMI data. c. Histograms representing the distribution of non-zero counts in raw UMI data across various data transformations. d. Histograms detailing the zero counts in raw UMI data, comparing VST with integrated data where zeros are imputed or converted to non-zeros. e. Histograms showing the distribution of gene RUNX3 across different data transformations.

## The curse of zeros

Bulk RNA-seq provides the average transcriptional output of each gene expressed within a population of heterogenous cell types^24, 25^. Even a moderate sequencing depth can yield information about many thousands of different genes. In comparison, scRNA-seq data is much sparser in comparison, with fewer genes expressed per sample and a high proportion of genes with zero UMI counts. Zeros in UMI counts for a gene can arise from three scenarios: a genuine zero, indicating that the gene is not expressed, or a sampled zero, indicating that the gene is expressed at a low level, or a technical zero, indicating that the gene is expressed at a high level but not captured by the assay. Despite an increasing body of evidence suggesting that cell-type heterogeneity is the major driver of zeros observed in 10X UMI data^26–28^, the prevailing notion within the single-cell community is that zeros are largely uninformative technical artifacts caused by “drop-out” genes (i.e., technical zeros).

Accordingly, many single-cell DE studies include pre-processing steps aimed at removing so-called zero inflation. Several popular pre-processing methods include: 1) performing feature selection by aggressively removing genes based on their zero detection rates, such as requiring non-zero values in at least 10% of total cells and restricting DE analysis to a smaller gene set; 2) imputing zeros and performing DE on imputed values^29–32^; or 3) modeling zeros explicitly as an extra component and essentially performing DE on non-zero values only^33, 34^.

However, if zeros are in fact biological zeros due to no expression or very low expression, dismissing or correcting for zeros in scRNA-seq is equivalent to discarding a significant portion of information in the dataset before any analysis. By failing to account for cell-type heterogeneity, zero-inflation pre-processing steps such as normalization and imputation become inappropriate and can introduce unwanted noise into downstream analyses, including DE. Ironically, the most desired markers in single-cell DE analysis—e.g., genes that are exclusively expressed in a rare cell type that accounts for less than 5% of the total population—may be obscured by current pre-processing steps for handling zeros.

In the fallopian tube dataset, we observed that distinct cell types display varied gene expression patterns in UMI counts. However, these differences become less apparent in imputed or certain transformed datasets (Fig. 1a). Gene expression frequency differs among cell types (Fig. 1b). However, normalization processes can substantially alter the distribution of both non-zero UMI (Fig 1c) and zero UMI counts (Fig. 1d) counts. For example, while the frequency of genes exponentially decline as raw UMI counts increase, VST data forms a more bell-shaped curve with a mode around 1.5 for non-zero raw UMI counts. Non-zero CPM-normalized data, (scaled by 1000) peaks near 0.2 and is more right-skewed than the VST data. Following batch integration, UMI counts primarily fall below 5 and are not as strongly right-skewed. It is noteworthy that zero UMI counts can be given non-zero values via normalization (except with CPM normalization); for example, zeros in VST data are adjusted to negative values and are left-skewed (Fig. 1d). Conversely, the integration process transforms original zeros to values clustered closely around zero. We further examined the distributions of gene expression from one gene. Using the gene RUNX3 as an example (Fig. 1e), the distributions in raw UMI counts and CPM data remain right-skewed. In contrast, the VST and integrated data showcase broader, bell-shaped distributions. The handling of zeros in these latter datasets (VST and integrated) intrinsically sets them apart from the former. This variability, combined with shifts in distribution skewness, may raise concerns when performing DE analysis with normalized values.

## The curse of donor effects

Recent reviews have highlighted that many single-cell DE analysis methods are susceptible to generating false discoveries^5^. This is mainly due to failing to account for variations between biological replicates, commonly referred to as “donor effects”. In single-cell studies, donor effects are always confounded with batch effects since cells from one biological sample are typically processed in the same experimental batch. While single-cell studies that contain multiple samples will perform batch correction as pre-processing, they usually do not correct for donor effects when performing DE tests in the downstream analysis.

One question that arises is whether batch effect correction alone suffices to eliminate donor-related effects. To address this, we investigated the contributions of variations from different sources before and after batch correction. Using the same fallopian tube dataset, we further separated 4553 T/NK cells into 20 subtypes using HIPPO^35^ (Fig. 2a, S1). With the aid of canonical markers, we identified specific subtypes, including NK, CD4+ T, CD8+ T and mature naive T cells. We then focused on subtypes that were observed in all donors (Fig. 2bc).

**Figure 2:**
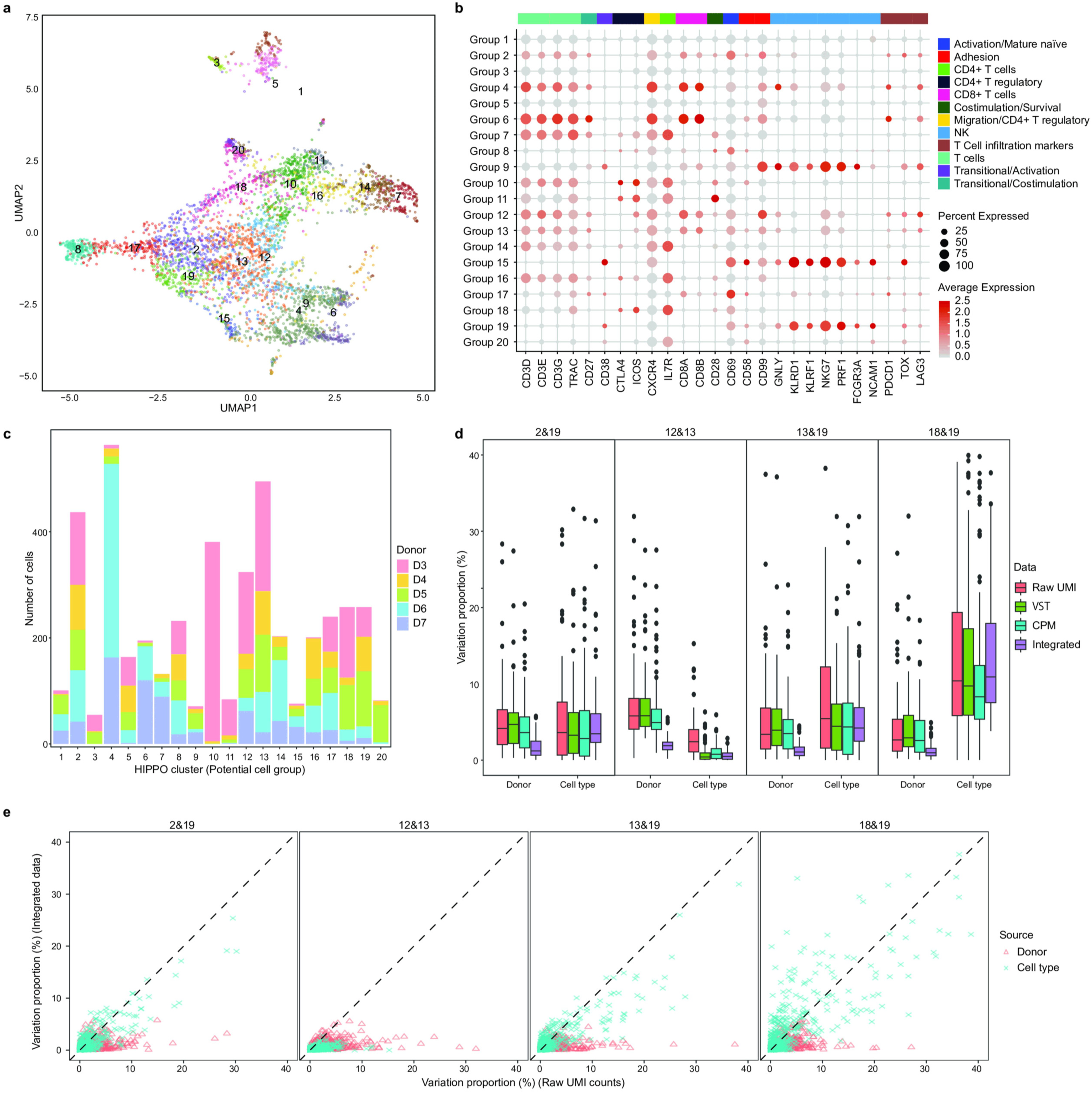
Cluster and Variation Analysis of Single-Cell Data from the Fallopian Tube in Case Study 1. a. UMAP visualizing 20 clusters identified by HIPPO in case study 1. b. Canonical markers delineate specific cell subtypes: clusters 9, 15, and 19 as NK cells; clusters 7, 10, 11, 14, 16, 18, and 20 as CD4+ T cells; clusters 4, 6, 12, and 13 as CD8+ T cells; clusters 8 and 17 as mature naive T cells. c. Distribution of donors across the 20 identified clusters. d. Comparative analysis of variation proportions attributable to donor and cell type effects across different pairs and datasets. e. Scatter plots comparing variation proportions due to donor and cell type effects across various pairings and data sources.

To quantify the proportion of variation originating from different sources, we fit a linear model, using cell types and donors as covariates, for each gene in several subtype pairs. Through all pairs, the integration led to a reduction in donor variation (Fig. 2d, S2). However, in comparisons of two subtypes of the same cell type (12 vs.13) and two subtypes of different cell types (13 vs. 19), we observed a decrease in the proportion of cell-type–related variation. This underscores that integration not only mitigates batch effects but also impacts the phenotypes of interest. Importantly, our analysis indicated that even after implementing batch correction, a notable percentage of genes still exhibited donor-related effects (Fig. 2e). As batch effects are often estimated from leading principal components, representing a consensus from a subset of genes, it is quite possible that residual donor effects persist on some, if not all, genes. Therefore, it is crucial to account for donor effects when performing DE tests to avoid false discoveries and obtain accurate results, even after removing batch effects.

One popular solution to address the issue of donor effects in single-cell studies is the use of pseudo-bulk analysis. This approach involves merging cells from the same donor and treating the resulting data as bulk RNA-seq. DE analysis is then performed using tools such as DESeq2^36^ or edgeR^37^. However, this framework ignores within-sample heterogeneity by treating donor effects as a fixed effect and assumes that each cell from the same donor is equally affected. As a result, this type of analysis can be overly conservative and potentially lead to missed discoveries^5^. Moreover, bulk RNA-seq DE tools typically perform normalization by default, which may have the same drawbacks mentioned earlier in the context of single-cell studies. Thus, caution is advised when using pseudo-bulk analysis as it may not always provide an accurate solution to the problem of donor effects in single-cell studies.

## The curse of cumulative biases

In scRNA-seq analysis, it is common to follow a hierarchical, sequential workflow for clustering and DE analysis. This approach can carry forward biases from one step to the next, from batch correction through to normalization, imputation, and feature selection. Such cumulative biases can ultimately diminish the power to detect differentially expressed genes.

Unsupervised learning, especially clustering analysis, is essential in single-cell studies. It groups cells based on gene expression patterns, facilitating the cell-type annotation. While clustering is effective with normalized values like CPMs, it essentially reweights gene features based on their relative contributions. As a result, clustering provides a generalized perspective of variation in gene expression across cell types. The reliance on relative expression also makes clustering fairly resilient to errors and biases introduced by the pre-processing steps.

On the other hand, DE analysis operates at the gene level, using group labels from the clustering process. The effects of biases, whether from donors or batch processing, can vary for each gene. Although DE analysis technically follows clustering—given its reliance on group labels—the metrics used do not need to be identical for both. As we show later in the case studies with data that complete clustering and annotation successfully, if DE analysis is performed using processed expression levels, the cumulative biases can still lead to false discoveries or overlook of certain DEs.

## An alternative paradigm – mixed effects model on UMI counts

To minimize the pre-processing biases discussed above, we proposed an approach that conducts DE analysis on raw UMI counts prior to implementing batch correction, normalization, imputation, or feature selection. This approach, which uses a generalized linear mixed model (GLMM)^38^, preserves sample-specific structures and biological signals in the data. Furthermore, our proposed approach can adjust for any potential confounding factors, such as batch, age, sex, or ancestry, by incorporating them as covariates with fixed effects. This framework enables us to explicitly account for the variation among biological replicates in comparison to other effects (Fig. 3). The proposed procedures have been implemented in software LEMUR (https://github.com/C-HW/LEMUR).

**Figure 3.**
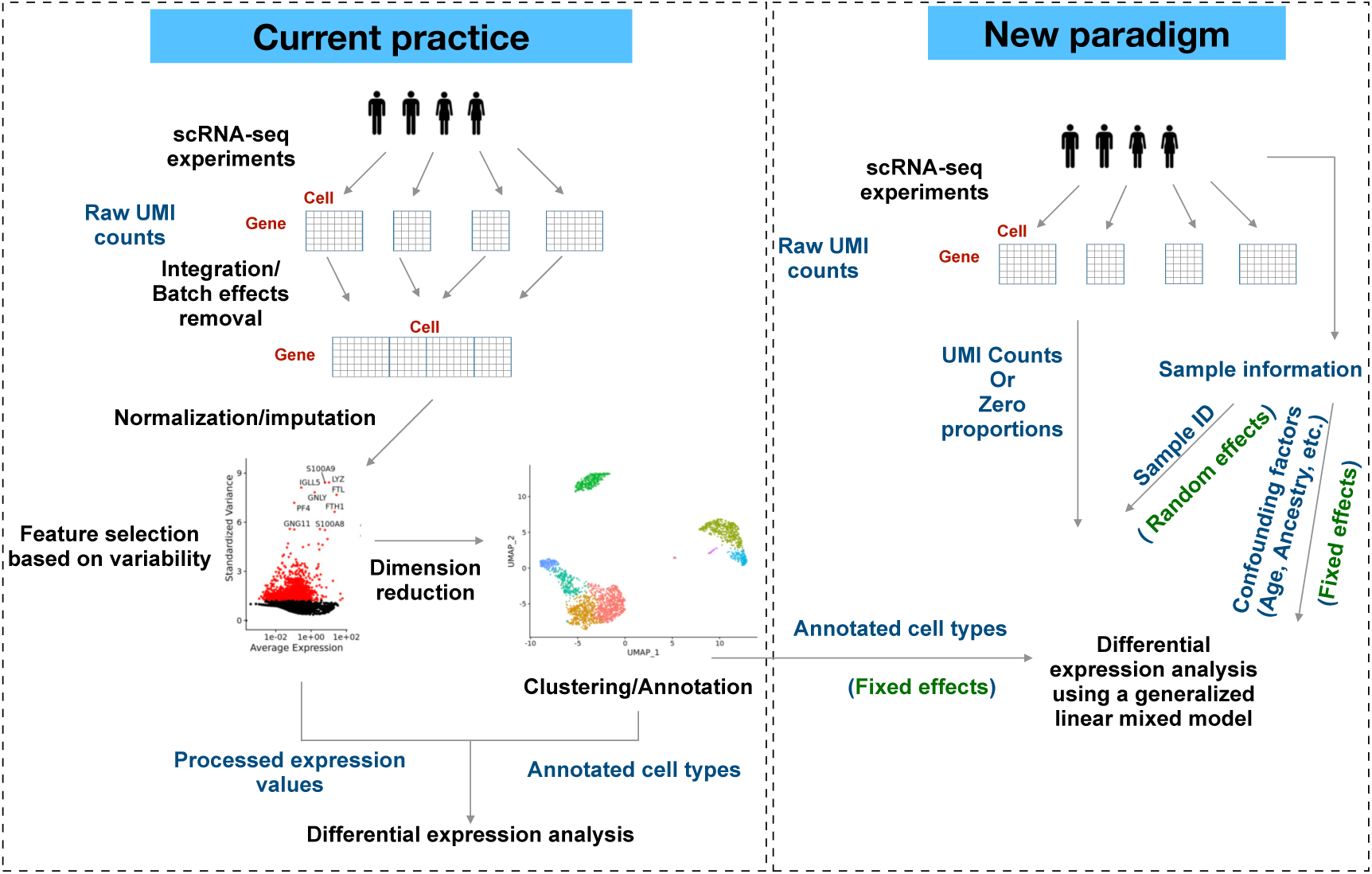
Comparison of established workflows and proposed paradigm for single-cell analysis. Left: Under the current single-cell analysis pipeline, the raw UMI counts collected from multiple donors are integrated to remove the batch effects and normalized for further analysis. It is common to perform DE analysis on processed data. Right: Our new paradigm directly performs a generalized linear mixed model on raw UMI counts. The random effect can account for the batch effect due to samples. The annotated cell types can be obtained from existing pipeline or HIPPO algorithm which clusters cells based on the zero proportions of UMI counts.

Unlike existing packages that utilize GLMMs, such as Muscat^39^, LEMUR treats group-of-interest as a fixed effect while accounting for donor-specific variations as random effects. In contrast, Muscat assigns a random effect term for each combination of donor and group-of-interest. Muscat’s approach treats certain aspects of group-of-interest variability as random effects, potentially masking differences between groups. Furthermore, Muscat’s GLMMs use library size as an offset to normalize counts, essentially focusing on relative abundance rather than raw counts. Overall, Muscat’s GLMMs operate similarly to pseudo-bulk methods, grouping counts within the same donor before performing the normalization, which can result in comparable performance, as demonstrated in the later examples.

To benchmark the performance of our new paradigm, we implemented eight distinct methods for DE analysis: two new paradigm methods, Poisson-glmm and Binomial-glmm; two traditional pseudo-bulk methods DESeq2 and edgeR; and four existing single-cell–specific methods, MAST^34^, Wilcox in Seurat, and two Muscat GLMMs (MMvst and MMpoisson).

Binomial-glmm fits a GLMM model on the zero proportion of each gene, adding donors as random effect. Pseudo-bulk DESeq2 applies both VST and library size normalization. EdgeR applies library size normalization. MAST adopts a zero-inflated negative binomial model, using log-transformed CPM counts and incorporating the cellular detection rate as covariates. The Wilcox test is non-parametric, using integrated normalized counts. The two Muscat models, MMvst with VST counts and MMpoisson with raw UMI counts, account for library size. Both Muscat models consider donor–group combinations as random effects. See ‘Methods’ for more details.

Each method was rigorously evaluated in two case studies (across cell types and across cell states) and under different scenarios, such as variations in library size between groups and pronounced heterogeneity within groups.

## Case study 1 – DE analysis on different immune cell types in fallopian tube

In this dataset, we examined the efficacy of various methods across three distinct scenarios: homogeneous groups with differing library sizes, homogeneous groups with similar library sizes, and heterogenous groups. For each scenario, we illustrate the overarching gene expression profile, describe the DE results using diagnostic plots, and conduct a gene ontology (GO) analysis to investigate the biological foundations of our DE findings.

### Contrasting CD8+ T cell subgroups with marked library size differences

The first comparison is between groups of CD8+ T cells (clusters 12 and 13), where there are notable differences in library sizes (Fig. 4a). This example illustrates the impact of library-size– based normalization on single-cell data. Using a two-sample t-test, we compared gene expression means between these groups with raw UMI counts and three other normalization methods (Fig. 4b) using absolute t-scores. While t-scores from CPM mirror those from UMI counts, albeit with minor shrinkage, both VST and integration show substantial shrinkage. This normalization process dampens the gene expression differences between the groups before deploying any DE detection techniques.

**Figure 4.**
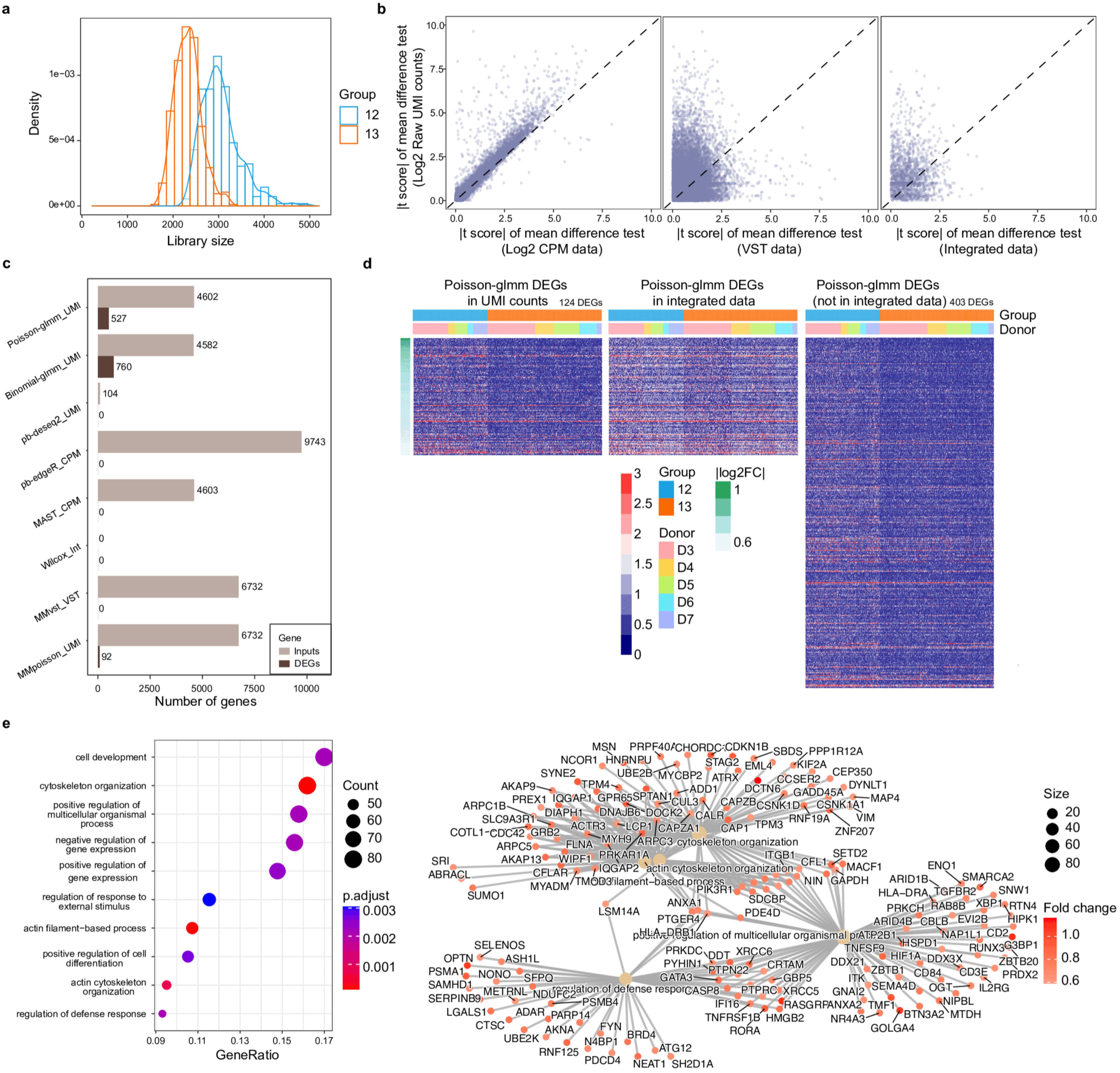
DE analyses on CD8+ T cell subgroups. a. Density plot of the library size for group 12 and 13. b. Scatterplot comparisons of t-scores from mean difference tests between raw UMI counts and other transformed data. Each gene’s expression in two different groups is compared, showcasing the pairwise absolute t-scores from various data sources. c. Counts of input genes and DEGs in different DE methods. d. Heatmaps visualize Poisson-glmm DEGs. Order: UMI counts (left), integrated data (middle), and genes not included in the integrated data but shown in UMI counts (right). Heatmaps arrange genes by descending Poisson-glmm fold change estimates and columns group cells by cell clusters and donors. e. GO analysis of the DEGs identified by Poisson-glmm.

Each method employs its unique filtering approach within the implemented function, resulting in varying numbers of input genes. Specifically, Poisson-glmm, Binomial-glmm, and MAST utilized nearly 4600 genes as input (Fig. 4c). In contrast, pseudo-bulk DESeq2 applied default quality control criteria to both genes and cells, resulting in only 104 genes being retained. Pseudo-bulk edgeR retained 9743 genes in the CPM data as inputs, while Muscat mixed models utilized 6732 genes. Notably, the Wilcox method from the Seurat package yielded no genes that passed the default filtering procedure. However, when a more lenient filtering criterion was applied, the impact on the differential expression results remained minimal.

In the volcano plots, both Poisson-glmm and Binomial-glmm display heavily imbalanced expression patterns, aligning with the observations in the density plots (Fig. S3b). However, the other methods do not reflect this observation, with fold change estimates appearing evenly spread. The histograms of adjusted p-values for other methods are concentrated in large values (Fig. S3c). Pseudo-bulk methods and mixed models from the Muscat package, in particular, exhibit p-values that are clustered around one. Despite observing imbalanced expression patterns in density plots and volcano plots in this comparison, only our GLMM methods identify a substantial number of differentially expressed genes (DEGs) (Fig. 4c). The heatmaps of DEGs further emphasize that raw counts can better capture the differences between groups compared to integrated counts (Fig. 4d). Furthermore, 403 DEGs were excluded from the integrated data before testing.

The DEGs prominently feature GO terms associated with actin cytoskeleton reorganization and immune synapse formation (Fig. 4e). As T cells detect antigens on an antigen-presenting cell, they establish an immunological synapse, necessitating substantial actin filament restructuring. Actin polymerization within this synapse aids the transit of receptors and signaling molecules, crucial for T cell activation. Our results hint that among these two CD8+ T cell groups, group 12 cells are actively recognizing antigens. Cell groups 12 and 13 had notable differences in library sizes. While the DEGs we identified contributed to the disparity in measured RNA content between the two groups, genes that were not differentially expressed had a much larger effect on library size; consequently, normalization erased the contribution of the DEGs to differences in expression patterns. Accordingly, in this example, only our GLMMs, which operate directly on UMI counts, successfully identified DEGs.

### A Glimpse at CD4+ T Cells vs. NK Cells: No Striking Library Size Differences

The second comparison is between CD4+ T cells and NK cells (clusters 2 and 19). In the density plot, we observed similar library sizes based on UMI counts for the two clusters across donors except for donor 7 (Fig. 5a). The zero-proportions of genes in these two clusters fit a Poisson distribution well, indicating relative homogeneity within each cell cluster (Fig. 5a).

**Figure 5.**
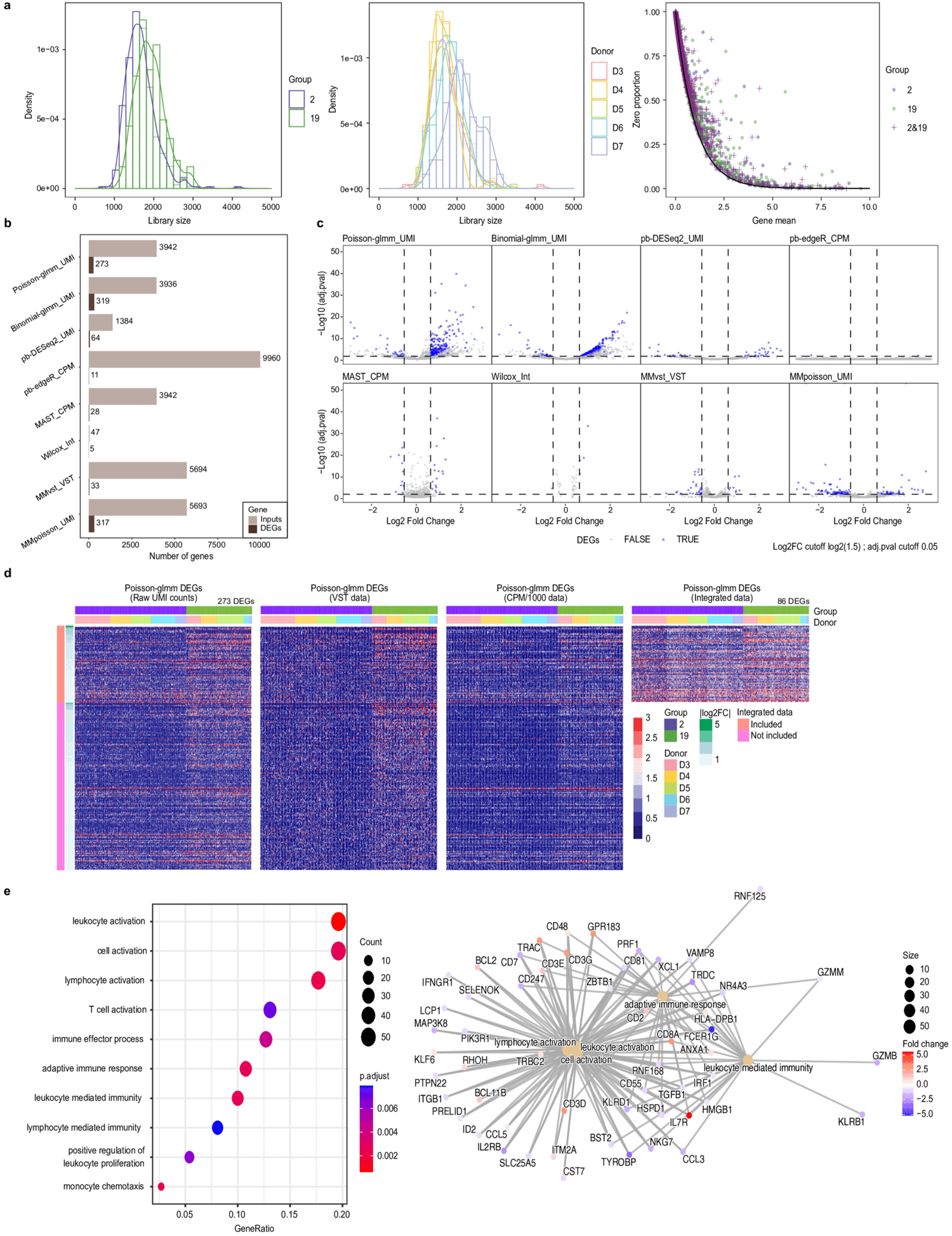
DE analyses on CD4+ T Cells vs. NK Cells. a. Left: Density plot of the library size for group 2 and 19. Middle: Density plot of the library size by different donors. Right: Zero proportion plots for each group and combined groups. b. Counts of input genes and DEGs across different differential expression methods. c. Volcano plots for each method, highlighting DEGs in blue. The signs of log2 fold change are adjusted such that positive signs represent higher expression in group 19. d. Heatmaps of Poisson-glmm DEGs shown in different data sources, with genes in integrated data featured in the top block, and those absent in the lower block. e. GO analysis of the DEGs identified by Poisson-glmm.

In this comparison, Poisson-glmm, Binomial-glmm, and MAST utilized nearly 4000 genes as input (Fig. 5b). Methods implemented in the Muscat package, including pseudo-bulk methods DESeq2 and edgeR, as well as mixed models MMvst and MMpoisson, employed 1384, 9960, 5694, 5693 genes, respectively, in accordance with their filtering procedure. Notably, the Wilcox method from the Seurat package includes only 47 genes as input due to the filtering based on the log2 fold change between two groups of interest. The log2 fold change in the package is calculated using the formula log2(1 + mean1) / log2(1 + mean2) on the input data, which can be normalized/integrated data by Seurat or other packages. This transformation attenuates the ratio of the two group means through the addition of 1 to each mean, resulting in the exclusion of a substantial number of genes. Wilcox, MAST, pseudo-bulk methods, and MMvst each identified fewer than 100 DEGs. In contrast, the methods that use UMI counts, Poisson-glmm, Binomial-glmm, and MMpoisson, identified 273, 319, and 317 DEGs, respectively (Fig. 5b).

In the volcano plots, there are more positive estimates of log2 fold change by Poisson-glmm and Binomial-glmm, signifying that genes are more expressed in cluster 19 (Fig. 5c). From the pairwise comparisons of log2 fold change (Fig. S4b), MAST, Wilcox, and MMvst exhibit smaller log2 fold change estimates, due to normalization processes that shrink the values. Pseudo-bulk methods tend to yield more conservative p-values (Fig. 5c, S4c), as illustrated in the histograms (Fig. S4d). While the log2 fold change estimates are consistent across our GLMMs, pseudo-bulk methods, and MMpoisson, the presence of deviant p-values leads to significant disparities in the identification of DEGs. Our GLMMs identified many more DEG candidates, surpassing the thresholds of adjusted p-value and fold change.

In Figure 5d, we display gene expression from DEGs identified by Poisson-glmm alongside heatmaps for VST, CPM, and integrated data. Notably, differences among these heatmaps are subtler than those displayed in raw UMI counts. The integrated data displays elevated gene expression across groups, obscuring distinctions. The heatmaps of DEGs from Poisson-glmm and Binomial-glmm show the validity of DEGs (Fig. S4f), while most of the DEGs identified by MMpoisson do not display any differential expression pattern in UMI counts (Fig. S4g). We performed gene ontology (GO) enrichment analysis on DEGs from Poisson-glmm. The DEGs are enriched for GO terms related to leukocyte activation, cell activation, and lymphocyte activation (Fig. 5e), suggesting NK cells represented by cluster 19 are more active than the CD4+ T cells.

In summary, in the comparison of two cell clusters of similar library sizes, normalization continued to obscure informative differences between the two clusters and hindered the identification of potential DEGs.

### Deciphering the Complexities of Heterogeneous Groups: Mature T Cells vs. CD4+ T Cells

Finally, by merging groups 8 and 17 and groups 2 and 19, we created two less homogenous groups-of-interest: mature T cells and CD4+ T cells, respectively. The distribution of library sizes between these clusters exhibits noticeable differences (Fig. 6a), and the zero proportions of these groups deviate from a Poisson distribution (Fig. S5a).

**Figure 6.**
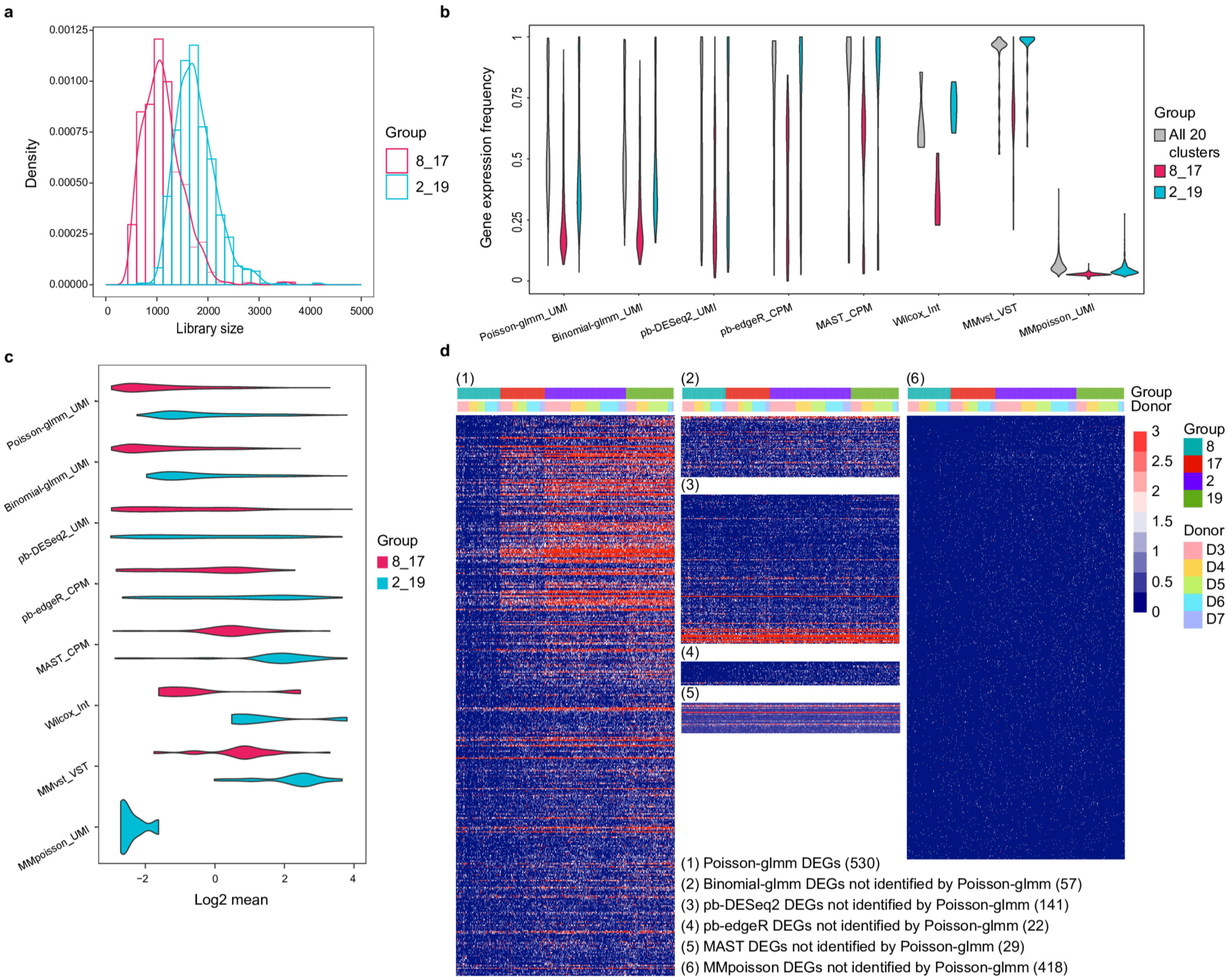
DE analyses on heterogeneous groups: Mature T Cells vs. CD4+ T Cells. a. Density plots comparing library sizes for combined groups 8 & 17 and 2 & 19. b. Comparisons of the gene expression frequency of the DEGs from different methods. c. Violin plot of log2 gene mean for DEGs from different methods. d. Heatmaps of DEGs from different methods.

In this comparison, Poisson-glmm, Binomial-glmm, and MAST used ∼3480 genes as input. Pseudo-bulk DEseq2, edgeR, and mixed models utilized 1937, 10483, 7099 genes, respectively. For Wilcox, 123 genes passed the filtering procedure. The volcano plots revealed similar patterns to previous comparisons across various methods (Fig. S5c). Our GLMM methods exhibited predominantly positive estimates of fold change, suggesting higher expression of abundant genes in CD4+ T cells (group 2&19). MAST, and MMvstn showed a somewhat similar tendency, but less imbalanced. However, pseudo-bulk methods and MMpoisson provided evenly distributed estimates in both directions.

The estimates of log2 fold change are not quite identical among different methods (Fig S5e). Both pseudo-bulk methods exhibited a negative shift compared to Poisson-glmm, while MMpoisson had a positive shift. MAST, Wilcox and MMvst showed shrinkage as before. Additionally, most input genes for the Wilcox method displayed positive fold changes, albeit with small magnitudes. This observation sheds light on how normalization and logarithmic transformation during pre-processing influences the estimation of differences in gene expression.

When we examine the violin plots of gene expression frequency and log2 mean for the DEGs identified by each method, it becomes apparent that MAST, Wilcox, and MMvst captured fewer DEGs with lower gene expression frequency and smaller gene means than the remaining methods (Fig. 6b, 6c). It is worth noting that MAST is a zero-inflated model, which incorporates excessive zeros as an additional component. However, MAST might not effectively characterize the zeros, as demonstrated in previous studies on UMI counts^26^. Consequently, potential DEGs that are lowly expressed may be masked by the model. The Wilcox method tends to filter out a substantial number of genes, which poses challenges in identifying lowly expressed genes. MMvst, despite having a considerable number of input genes (n=7099), only identified 35 DEGs.

The heatmap of DEGs in Poisson-glmm reveals distinct expression patterns between the two groups (Fig. 6d (1)). However, in this example, the inherent heterogeneity within each group impacts the fitness of Poisson model, potentially leading to false discoveries. To evaluate the possibility of false discoveries by Possion-glmm, we examined DEGs identified by other methods, but not by Poisson-glmm (Fig. 6d (2)-(6)). The heatmaps make it evident the DEGs that differentiate between the two groups are largely identified by Poisson-glmm only; the other methods did not contribute additional valid DEGs that differentiate the two groups. Conversely, most of the DEGs detected by Poisson-glmm exhibit differential expression despite the heterogeneity within each group.

Notably, MMpoisson mainly detected DEGs with small means (Fig. 6c), not showing clear differences between different groups (Fig. 6d (6)). And the DEGs are mutually exclusive to those identified by Poisson-glmm. Although Poisson-glmm and MMpoisson both use UMI counts, MMpoisson includes group information as a random variable and involves library size as an offset; our result underscores the significance of using an appropriate random effect in a mixed model and suggests that the cell group information should be excluded from the random component.

The DEGs are enriched for GO terms related to peptide metabolic process and cytoplasmic translation, indicating lower ribosomal RNA activities in mature T cells (Fig. S5g). Indeed, mature T cells exhibit lower levels of ribosomal RNA activity compared to their immature counterparts, mainly due to the state of activation and the metabolic requirements of the cells. On the other hand, mature T cells, which are not rapidly proliferating, have less need for protein synthesis and thus exhibit lower levels of rRNA activity. However, upon antigen recognition and activation, mature T cells can rapidly upregulate rRNA activity and protein synthesis to support clonal expansion and effector function. This differential regulation of rRNA activity is one of the many ways in which cells regulate their metabolic activities to adapt to different physiological conditions.

In this example, Poisson-glmm detected more valid DEGs for heterogenous cell populations than other methods. Normalization still diminished measurable differences between groups. We also raise concerns about the masking of lowly expressed genes by the improper treatment of zeros, as seen in MAST method and VST data.

## Case study 2 – DE analysis on different states of B cells

In this case study, we applied our proposed DE framework to data collected by Kang et al^40^; this dataset consists of 29,065 cells and 7,661 genes from eight distinct cell types, collected from peripheral blood mononuclear cells of eight lupus patients. Within each cell type, the cells are evenly split into two groups for perturbation: unstimulated control and IFN-β stimulated (Fig. S6a). UMAP plots (Fig. 7a) highlight that gene expression patterns are more differentiated between stimulation states than between cell types. The zero-proportion plots fit better to Poisson distribution when separated by stimulation states than only by cell types (Fig. S6b). This observation motivated us to focus on DEGs between the cell states rather than between the cell types.

**Figure 7.**
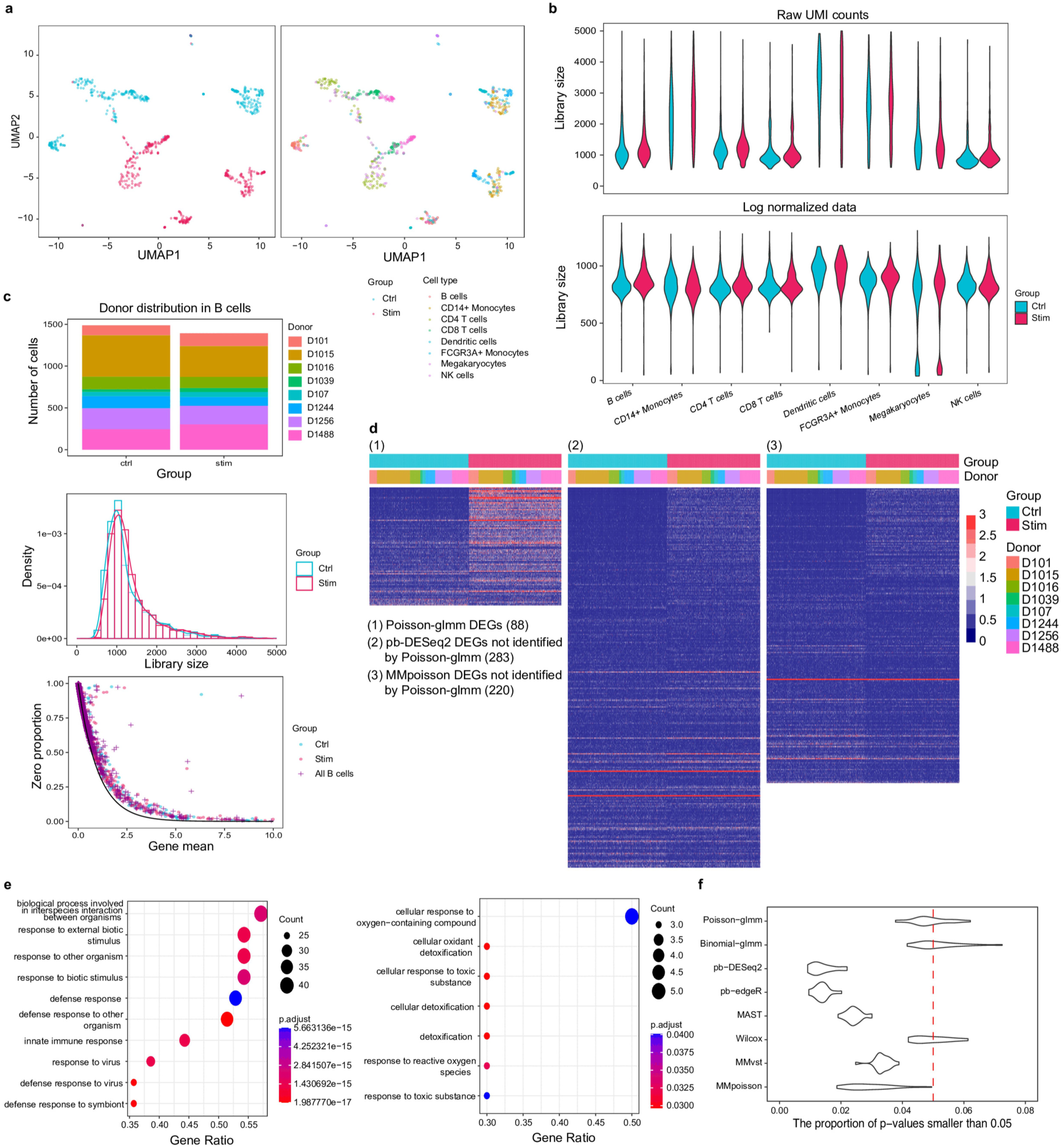
Overview of case study 2 and DE analyses on different states in B cells. a. UMAP showing groups and cell types for case study 2. b. Library size comparisons before (raw UMI counts) and after normalization (log-normalized data) by cell type. c. Top: Donor distribution among B cells. Middle: Density plot of library size in different states. Bottom: Zero proportion plots for different states and combined states. d. Heatmaps of DEGs identified from different methods. e. Violin plots depicting the proportion of p-values below 0.05 for each method. f. GO analysis for up-regulated (left) and down-regulated genes (right).

Like the previous case study, we found that the distribution of library sizes underwent significant changes after normalization (Fig. 7b). Raw UMI counts show that each cell type has a unique library size distribution. However, these differences became less pronounced following normalization, while library sizes remained relatively consistent between states within a single cell type. Normalization seems to predominantly affect differences across cell types rather than between states.

For the remainder of our case study, we focused on B cells. The cells from each donor were divided approximately equally between the control and stimulated groups (Fig. 7c top), and the library size distribution in these two groups is similar (Fig. 7c middle). The zero-proportion plot suggests that the data does not perfectly fit the expected curve from the Poisson distribution, indicating the presence of a mixture of subtypes within B cells (Fig. 7c bottom).

In our analysis of the subset comprising unstimulated and stimulated B cells, the majority of DE methodologies used about 2,550 genes as inputs (Fig. S7a). However, the Wilcox approach within Seurat selected only 144 genes. The estimates of fold change for the two states in B cells exhibit an even spread across all methods, as depicted in the volcano plots (Fig. S7b). MAST and MMvst struggled to identify differential patterns. Different from previous examples, our GLMM approach flagged fewer DEGs than both pseudo-bulk techniques and MMpoisson. Notably, the DEGs that were not shared between pseudo-bulk DESeq2 or MMpoisson and Poisson-glmm predominantly belong to the extremely low expression category (Fig. 7d (2), (3)).

We hypothesized that this result could be explained by using fold change as a DEG criterion. In bulk RNA-seq, a gene is typically labeled as a DEG if its adjusted p-value is below a certain threshold, often 0.05, and the fold-change estimate exceeds a predetermined value, typically 1.5 or 2 (Fig. S8a). Most single-cell DE methods use the same criteria. However, in single-cell datasets, the mean counts for many genes are exceedingly close to zero. Consequently, fold change may not be a reliable metric to differentiate nuances in read counts. For instance, if gene means are 2^-3^ for one group and 3^-3^ for another, the fold-change threshold of 1.5 is met, but the actual difference is a mere 0.0625, which does not convey a significant disparity in expression, especially when juxtaposed with genes having larger means. Moreover, near-zero values can result in computational inaccuracies, causing ratio deviated from the underlying true value.

To overcome the limitation of using fold-change ratios on small counts, we established a new criterion for DEGs based on absolute differences. Specifically, we mandated that the mean difference between two groups exceeds a set threshold, such as −1. In the volcano plot, numerous genes would be designated as DEGs when relying on ratio-defined fold change. Yet, as shown in the mean vs. mean difference plot that many genes that meet the p-value criteria showcase only modest changes in absolute means (Fig. S9a). This approach emphasizes genes with significant absolute differences, yielding more biologically pertinent results.

We performed GO enrichment analysis on up-regulated and down-regulated genes separately (Fig. 7e). We found IFN-β stimulated B cells have increased activities in interaction between organisms, defense response, defense response to virus and defense response to symbiont, while their activities in translation and other metabolic processes are decreased. Pseudo-bulk technique detected similar GO terms while MMpoisson detected very different down-regulated GOs (Fig. S9).

In this example, we demonstrated that conventional metrics to detect DEGs, especially fold change based on ratios, are ill-suited for low-count data where the large fold changes reported by current methods may be attributed to the ratio of two very small gene means. Careful post-processing is needed to prioritize signals and manage false discoveries.

### False discovery rates assessed under the null setting using permutation analysis

To assess p-value calibration in empirical data, a permutation analysis was conducted within a null dataset focusing on a group of interest. We specifically conducted the analysis on three datasets: the control group of B cells, group 2, and group 13 in case study 1. Each underwent random assignment to either the control or stimulus group. Subsequently, p-values for each gene were computed employing various methods, with the gene set confined to those input into the Poisson-glmm model. To mitigate potential gene filtering, the threshold for the Wilcox method was relaxed. This process was iterated 20 times, and on each iteration the proportion of p-values below 0.05 was calculated along with the corresponding false discovery of differentially expressed genes.

The analysis of the violin plot (Fig. 7f, S10) reveals that both our GLMM methods and the Wilcox method exhibit consistently well-calibrated p-values among different choices of null datasets. However, pseudo-bulk methods, and mixed models from Muscat appear excessively conservative, with an overall proportion considerably below 0.05. The performance of MAST is conservative in B cells but not in case study 1. The histograms of p-values across the 20 runs demonstrate a consistently flat distribution for our glmm methods and the Wilcox method, indicative of adherence to the null setting (Fig. S10). Conversely, other methods display overestimated p-values, yielding conservative outcomes. Note that even though Wilcox performed well in the permutation analysis, it is not powerful to detect real DEGs as shown in previous case studies. Under both the existing criteria and our newly established criteria for determining DEGs, each method detected, at most, one false discovery in each run.

## Discussion

In this manuscript, we examined existing DE approaches to pre-processing, input values and test statistics, and fold-change definitions in the context of single-cell DE analysis. We demonstrated through extensive real-data examples the limitations and drawbacks of current practices. We showed that current normalization and pre-processing techniques may obscure DEGs by an overreliance on relative RNA abundance and ignoring or correcting for biological zeros. We also illustrated how use of volcano plots in DE analysis, which also depends on relative RNA abundance, leads to false discoveries in lowly expressed genes by prioritizing fold changes in expression over absolute changes. We also argued that single-cell DE analysis suffers from false discoveries due to the inappropriate handling of donor effects, as well as from biases that accumulate as the consequence of sequential workflows.

We advocate a new paradigm, Poisson-glmm, which uses UMI counts as input and a generalized Poisson mixed effect models to account for batch effects and within-sample variation. This framework’s use of UMI counts can significantly improve current practices by leveraging absolute RNA expression. Poisson-glmm shows superior sensitivity and robustness toward model misspecification when compared to current single-cell DE methods, which should ultimately lead to new biological insights from single-cell data.

The use of UMI counts for DE analysis in scRNA-seq can significantly improve current practices, potentially making some current practices (e.g., volcano plots as a diagnostic DE tool) obsolete. However, relying on UMI counts as a representation of genuine RNA content predicates that measurements are strictly single-cell based, underscoring the need for meticulous doublet and triplet removal prior to DE analysis. Furthermore, seamlessly implementing this new paradigm into existing popular tools remains a challenge. Given this significant shift from current practices, a sustained effort will be required to educate and train researchers on these new alternatives and to reshape existing practices accordingly.

## Methods and materials

### Datasets and pre-processing

In case study 1, a 10X scRNA-seq dataset of post-menopausal fallopian tubes, with 57,182 cells sourced from five donors, covering 29,382 genes was analyzed. We obtained 20 clusters via HIPPO algorithm. We did not apply a pre-filtering procedure on this dataset, except for built-in filtering steps in each method. We used sctransform to get the VST data and the integration workflow provided by Seurat to obtain the integrated data.

All integration or normalization processes were performed on the entire dataset, since cell types are typically unknown during the pre-processing stage. In cross-batch integration, only the top 2,000 highly expressed genes were retained, which significantly reduced the number of genes for downstream analysis. The dataset had already been fully analyzed and annotated with cell types. We utilized the annotations to examine the effects of normalization/integration on distributions of library sizes across cells.

In case study 2, the dataset comprised 10X droplet-based scRNA-seq PBCM data from eight Lupus patients obtained before and after 6h-treatment with IFN-β. After removing undetected and lowly expressed genes (less than 10 cells expressing more than 1), the dataset consisted of 29065 cells and 7661 genes. The integrated data was replaced by log2-transformed normalized expression values obtained via computeLibrarayFactors and logNormCounts functions in Muscat.

### Poisson-glmm and Binomial-glmm

By default, we excluded any genes detected in fewer than 5% cells in the compared groups from differential testing. The GLMMs were implemented with glmmPQL function of the MASS package. We calculated adjusted p-values by using Benjamini-Hochberg correction. Each model fitting was applied on one gene and the two compared groups.

We fit Poisson-glmm on UMI counts. Each count *X*_*cgk*_ sampled from cell *c*, donor *k*, and gene *g*, was modeled by

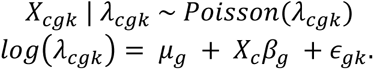

We fit Binomial-glmm on the zero proportions. Each count *X*_*cgk*_ was modeled by

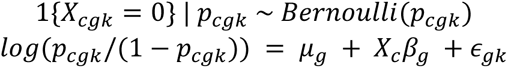

where *X*_*c*_ is the indicator for groups (e.g. cell types in case study 1, control/stimulus in case study 2), and *∈*_*gk*_∼*N*(0, *σ*_*g*_^2^) represents the random effects for donor *k*. Our goal was to test *H*_0_: *β*_*g*_ = 0.

For both methods, we provided “log2 fold change” computed by log_2_(exp(*β*_*g*_)). In Poisson-glmm, this estimate indicates the increment of log_2_(*λ*_2_) against log_2_(*λ*_1_), which is the conventional log2 fold change. However, this term in Binomial-glmm doesn’t represent the same meaning. It is the difference between logit(*p*_2_) and logit(*p*_1_). The p-value and BH adjusted p-value are provided.

### Benchmarked methods

Pseudo-bulk DESeq2 and pseudo-bulk edgeR are aggregation-based methods used in our comparison. The input counts were summed up for a given gene over all cells in each group and by donor. The pseudo-bulk data matrix has dimensions GxS, where S denotes the number of interactions of donors and groups. For example, if there are two groups and ‘a’ and ‘b’ donors in each group, then ‘S’ is equal to 2(a + b). We used raw counts as the input for DESeq2, while CPM counts were used for edgeR. The log fold change was converted to log2 fold change in all the comparisons. We implemented these two pseudo-bulk methods following the guided tutorial in Muscat package; https://www.bioconductor.org/packages/devel/bioc/vignettes/muscat/inst/doc/analysis.html.

For MAST, we fitted a zero-inflated regression model (function zlm) for each gene and applied a likelihood ratio test (function lrTest) to test for between-group differences in gene expression. Besides the labels of groups and the cellular detection rate, we also included donor labels in the covariates. This method was run on log (CPM+1) counts. We followed the tutorial https://github.com/RGLab/MAST.

Wilcox, a rank sum test, is the default DE method in the FindMarkers function in the Seurat package. We used integrated data and log counts as input. We computed the log fold change given in the output as log(1+mean1)/(1+mean2). We applied the default filter in FindMarkers to only test genes with a log fold change greater than 0.25. We calculated the adjusted p-value provided from the function based on Bonferroni correction. We followed the guided tutorial found here: https://satijalab.org/seurat/articles/de_vignette.

MMvst and MMpoisson are mixed models implemented in the Muscat package. MMvst fits linear mixed models on variance-stabilizing transformation data. MMpoisson fits Poisson generalized linear mixed models with an offset equal to the library size factors. In both models, we fit a ∼1 + group + (1∣sample) model for each gene, where ‘sample’ denotes the experimental units (the interaction of donors and groups). We followed the tutorial found at: https://www.bioconductor.org/packages/devel/bioc/vignettes/muscat/inst/doc/analysis.html.

### The criteria to determine DEGs

For the benchmarked methods, we adhered to conventional criteria for the identification of Differentially Expressed Genes (DEGs). Specifically, a gene was classified as a DEG if its absolute log2 fold change exceeded a predefined threshold, and the adjusted p-value was below a specified cutoff. Typically, DEGs are visually represented in volcano plots. In the first dataset, the log2 fold change threshold was set at log2(1.5), whereas in the second dataset, it was set at 1. The adjusted p-value threshold for both datasets was established at 0.05.

We proposed new criteria that are based on the convention plus the gene mean and the difference in mean. If the log2 gene mean in two groups is lower than a certain value (−2.25 in case study 1) and the log2 mean difference is smaller than a threshold (−1 in case study 1), the gene would not be considered as a DEG. These can also be used as a filter before any DE analysis to speed up the computation. Both criteria are adjustable, depending on the dataset’s performance and characteristics. An examination of heatmaps and mean difference against mean plot in advanced can be helpful to determine the thresholds when analyzing a new dataset (Fig. S8b, c).

### Variation analysis

To gain a deeper understanding of the donor effect and cell type effect concerning various types of counts, we conducted a variation analysis across multiple group comparisons. To ensure the consistency of our results, we restricted our analysis to genes presented in all datasets. For each gene, we employed linear models (lm (count ∼ donor + group)) and computed the variances attributed to three components: donor, group, and the residual. Logarithm transformation was applied to UMI counts and CPM data to address skewness. The outcomes of this analysis were then presented and compared based on the proportion of variation explained by the first two components across different count types and various pairs. The results of the top 500 genes with the lowest residual variations were exhibited.

### GO enrichment analysis

GO over-representation analyses were performed using the enrichGO function in the R package clusterProfiler with default parameters and the functional category for enrichment analysis set to the GO ‘Biological Processes’ category.

## Data availability

Both scRNA-seq datasets used in this study are publicly available. Processed and de-identified human single-cell RNA sequencing data scRNA-seq dataset of post-menopausal fallopian tubes has been deposited at Cellxgene under the following URL: https://cellxgene.cziscience.com/collections/d36ca85c-3e8b-444c-ba3e-a645040c6185. The droplet scRNA-seq data used in case study 2 is deposited under the Gene Expression Omnibus under the accession number GSE96583. The dataset is also available in R through the Bioconductor ExperimentHub package.

## Code availability

We provide an R package, LEMUR, implementing Poisson-glmm and Binomial-glmm methods for DE analysis discussed in this study. The LEMUR package is available from GitHub (https://github.com/C-HW/LEMUR). In addition, the R source code to reproduce all data analysis in the study is available from GitHub at https://c-hw.github.io/DEanalysis/index.html.

## Acknowledgements

The work was supported by National Institutes of Health grant R01 GM126553, R01 HG011883 and HG012927, and additional grant no. NSF 2016307 and Sloan Research Fellowship to M.C.

## Contributions

M.C. conceived and led this work. C.W. and M.C. developed the methods and performed the analyses. C.W. implemented the software. X.Z. participated in critically revising the draft. C.W. and M.C. wrote the paper with feedback from X.Z. All authors read and approved the final manuscript.

## Ethics declarations

Ethics approval is not applicable to this study.

## Competing interests

The authors declare no competing interests.

---

**Table 1.**
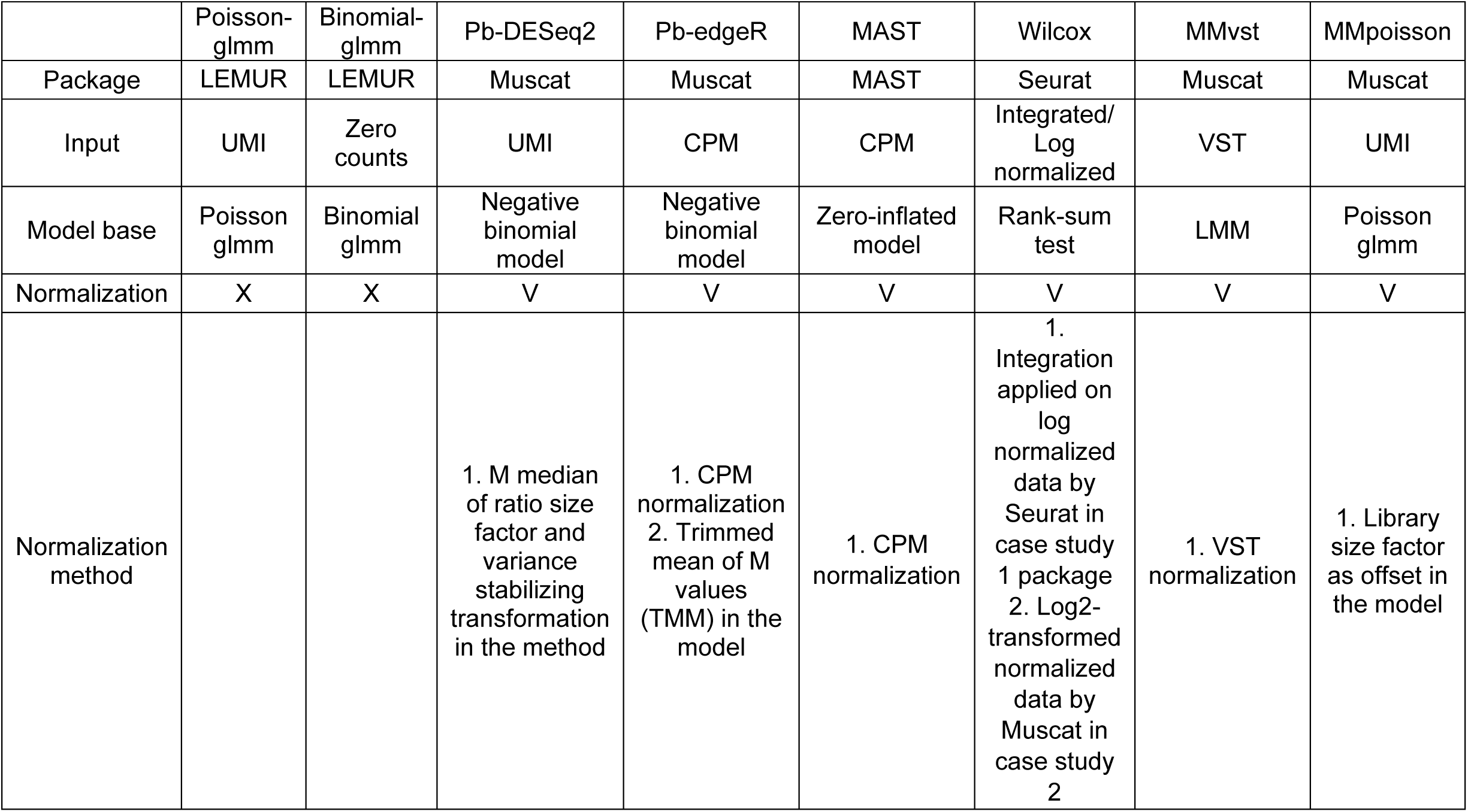
Comparison of DE methods used in this paper.

## Notes

https://c-hw.github.io/DEanalysis/index.html

### Competing Interest Statement

The authors have declared no competing interest.

## Reference

1. Saliba, A.-E., Westermann, A.J., Gorski, S.A. & Vogel, J. Single-cell RNA-seq: advances and future challenges. Nucleic acids research 42, 8845–8860 (2014).

2. Greenwald, W.W. et al. Pancreatic islet chromatin accessibility and conformation reveals distal enhancer networks of type 2 diabetes risk. Nature communications 10, 2078 (2019).

3. Grubman, A. et al. A single-cell atlas of entorhinal cortex from individuals with Alzheimer’s disease reveals cell-type-specific gene expression regulation. Nature neuroscience 22, 2087–2097 (2019).

4. Lawlor, N. et al. Single-cell transcriptomes identify human islet cell signatures and reveal cell-type–specific expression changes in type 2 diabetes. Genome research 27, 208–222 (2017).

5. Squair, J.W. et al. Confronting false discoveries in single-cell differential expression. Nature communications 12, 1–15 (2021).

6. Das, S., Rai, A., Merchant, M.L., Cave, M.C. & Rai, S.N. A comprehensive survey of statistical approaches for differential expression analysis in single-cell RNA sequencing studies. Genes 12, 1947 (2021).

7. Das, S., Rai, A. & Rai, S.N. Differential Expression Analysis of Single-Cell RNA-Seq Data: Current Statistical Approaches and Outstanding Challenges. Entropy 24, 995 (2022).

8. Lengyel, E., et al. A molecular atlas of the human postmenopausal fallopian tube and ovary from single-cell RNA and ATAC sequencing. Cell Reports 41 (2022).

9. Li, P., Piao, Y., Shon, H.S. & Ryu, K.H. Comparing the normalization methods for the differential analysis of Illumina high-throughput RNA-Seq data. BMC bioinformatics 16, 1–9 (2015).

10. Zyprych-Walczak, J. et al. The impact of normalization methods on RNA-Seq data analysis. BioMed research international 2015 (2015).

11. Dillies, M.-A. et al. A comprehensive evaluation of normalization methods for Illumina high-throughput RNA sequencing data analysis. Briefings in bioinformatics 14, 671–683 (2013).

12. Robinson, M.D. & Oshlack, A. A scaling normalization method for differential expression analysis of RNA-seq data. Genome biology 11, 1–9 (2010).

13. Lytal, N., Ran, D. & An, L. Normalization methods on single-cell RNA-seq data: an empirical survey. Frontiers in genetics 11, 41 (2020).

14. Leek, J.T. et al. Tackling the widespread and critical impact of batch effects in high-throughput data. Nature Reviews Genetics 11, 733–739 (2010).

15. Korsunsky, I. et al. Fast, sensitive and accurate integration of single-cell data with Harmony. Nature methods 16, 1289–1296 (2019).

16. Chen, M. & Zhou, X. Controlling for Confounding Effects in Single Cell RNA Sequencing Studies Using both Control and Target Genes. Sci Rep 7, 13587 (2017).

17. Chen, M. et al. Alignment of single-cell RNA-seq samples without overcorrection using kernel density matching. Genome research 31, 698–712 (2021).

18. Hu, J., Chen, M. & Zhou, X. Effective and scalable single-cell data alignment with non-linear canonical correlation analysis. Nucleic Acids Research (2021).

19. Schmid, R. et al. Comparison of normalization methods for Illumina BeadChip HumanHT-12 v3. BMC genomics 11, 1–17 (2010).

20. Tran, H.T.N. et al. A benchmark of batch-effect correction methods for single-cell RNA sequencing data. Genome biology 21, 1–32 (2020).

21. Hafemeister, C. & Satija, R. Normalization and variance stabilization of single-cell RNA-seq data using regularized negative binomial regression. Genome biology 20, 1–15 (2019).

22. Lause, J., Berens, P. & Kobak, D. Analytic Pearson residuals for normalization of single-cell RNA-seq UMI data. Genome biology 22, 1–20 (2021).

23. Argelaguet, R. et al. MOFA+: a statistical framework for comprehensive integration of multi-modal single-cell data. Genome biology 21, 1–17 (2020).

24. Wang, X., Park, J., Susztak, K., Zhang, N.R. & Li, M. Bulk tissue cell type deconvolution with multi-subject single-cell expression reference. Nature communications 10, 380 (2019).

25. Yang, Y. et al. Dimensionality reduction by UMAP reinforces sample heterogeneity analysis in bulk transcriptomic data. Cell reports 36 (2021).

26. Kim, T.H., Zhou, X. & Chen, M. Demystifying “drop-outs” in single-cell UMI data. Genome biology 21, 196 (2020).

27. Qiu, P. Embracing the dropouts in single-cell RNA-seq analysis. Nature communications 11, 1169 (2020).

28. Svensson, V. Droplet scRNA-seq is not zero-inflated. Nature Biotechnology 38, 147–150 (2020).

29. Gong, W., Kwak, I.-Y., Pota, P., Koyano-Nakagawa, N. & Garry, D.J. DrImpute: imputing dropout events in single cell RNA sequencing data. BMC bioinformatics 19, 1–10 (2018).

30. Li, W.V. & Li, J.J. An accurate and robust imputation method scImpute for single-cell RNA-seq data. Nature communications 9, 997 (2018).

31. Tracy, S., Yuan, G.-C. & Dries, R. RESCUE: imputing dropout events in single-cell RNA-sequencing data. BMC bioinformatics 20, 1–11 (2019).

32. Chen, M. & Zhou, X. VIPER: variability-preserving imputation for accurate gene expression recovery in single-cell RNA sequencing studies. Genome Biol 19, 196 (2018).

33. Pierson, E. & Yau, C. ZIFA: Dimensionality reduction for zero-inflated single-cell gene expression analysis. Genome biology 16, 1–10 (2015).

34. Finak, G. et al. MAST: a flexible statistical framework for assessing transcriptional changes and characterizing heterogeneity in single-cell RNA sequencing data. Genome biology 16, 1–13 (2015).

35. Kim, T.H., Zhou, X. & Chen, M. Demystifying “drop-outs” in single-cell UMI data. Genome Biol 21, 196 (2020).

36. Love, M.I., Huber, W. & Anders, S. Moderated estimation of fold change and dispersion for RNA-seq data with DESeq2. Genome biology 15, 1–21 (2014).

37. Robinson, M.D., McCarthy, D.J. & Smyth, G.K. edgeR: a Bioconductor package for differential expression analysis of digital gene expression data. bioinformatics 26, 139–140 (2010).

38. Clayton, D.G. Generalized linear mixed models. Markov chain Monte Carlo in practice 1, 275–302 (1996).

39. Crowell, H.L. et al. Muscat detects subpopulation-specific state transitions from multi-sample multi-condition single-cell transcriptomics data. Nature communications 11, 6077 (2020).

40. Kang, H.M. et al. Multiplexed droplet single-cell RNA-sequencing using natural genetic variation. Nature biotechnology 36, 89–94 (2018).

