## Supplementary Figures for "The curses of performing differential expression analysis using single-cell data"

**Supplementary materials for "The curses of performing differential expression analysis using single-cell data"**

### Supplementary Figures

#### Figure S1

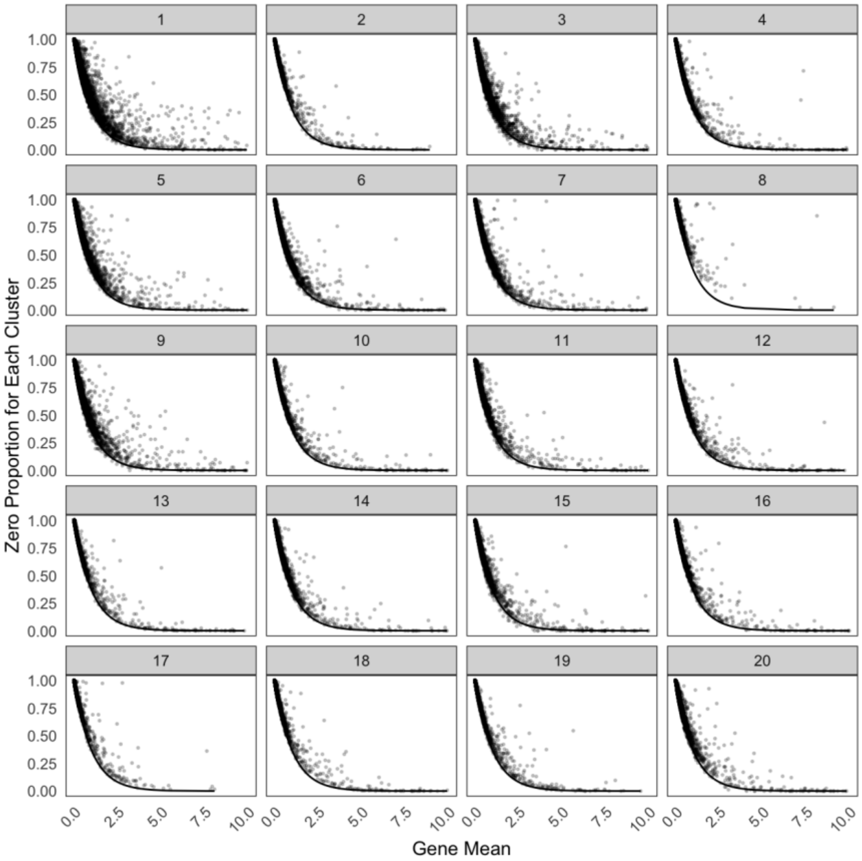


**Figure S1:** Zero proportion plots for each cluster obtained by HIPPO for case study 1.

### Figure S2

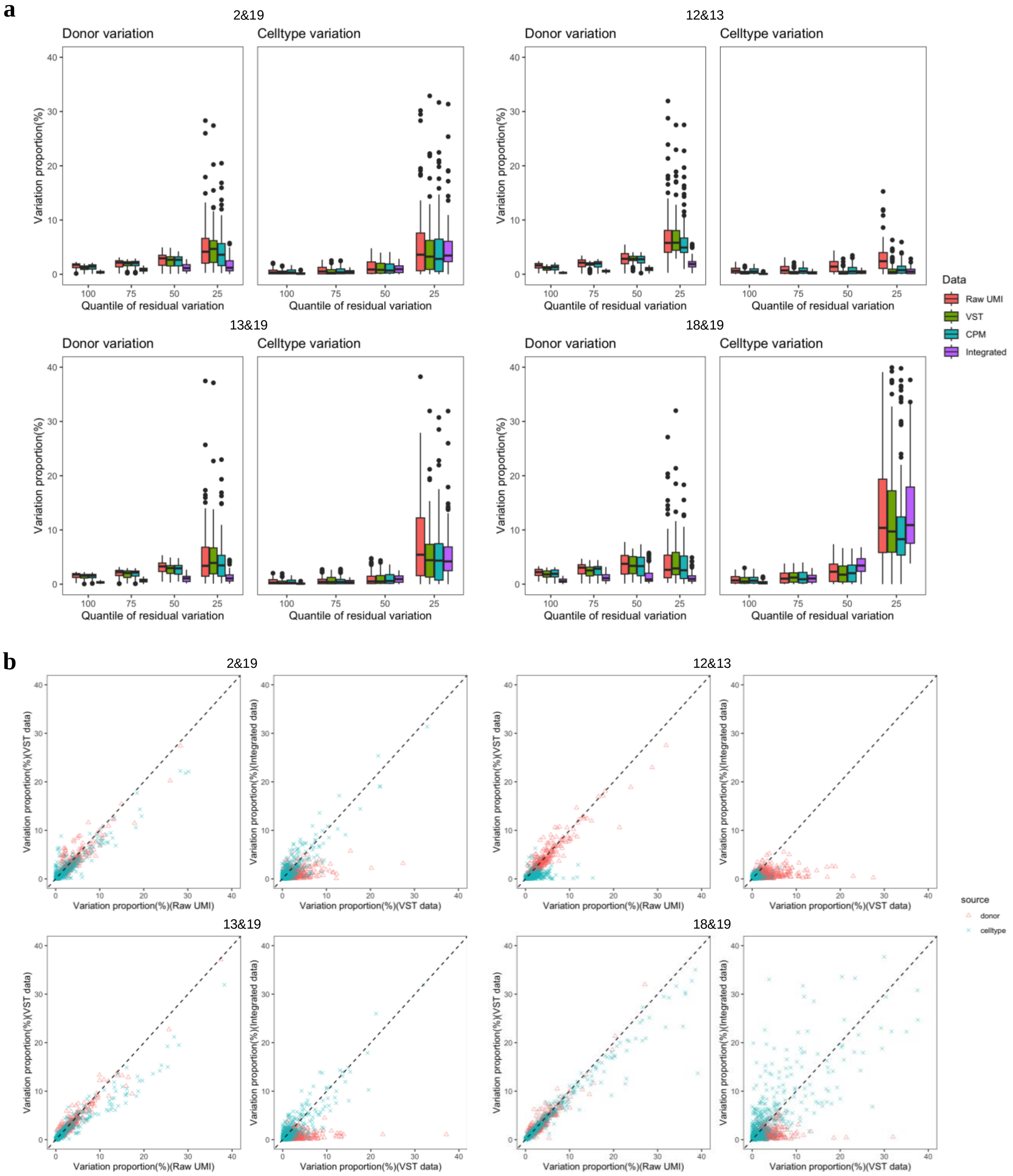


**Figure S2:** Additional variation proportion analysis results. **a)** Boxplots of donor variation and cell type variation grouped by quartiles of residual variation, displayed in different pairs and different data sources. **b)** Scatter plots of variation proportions for donor effects and cell type effects in different pairs and different data sources.

**Figure S3**

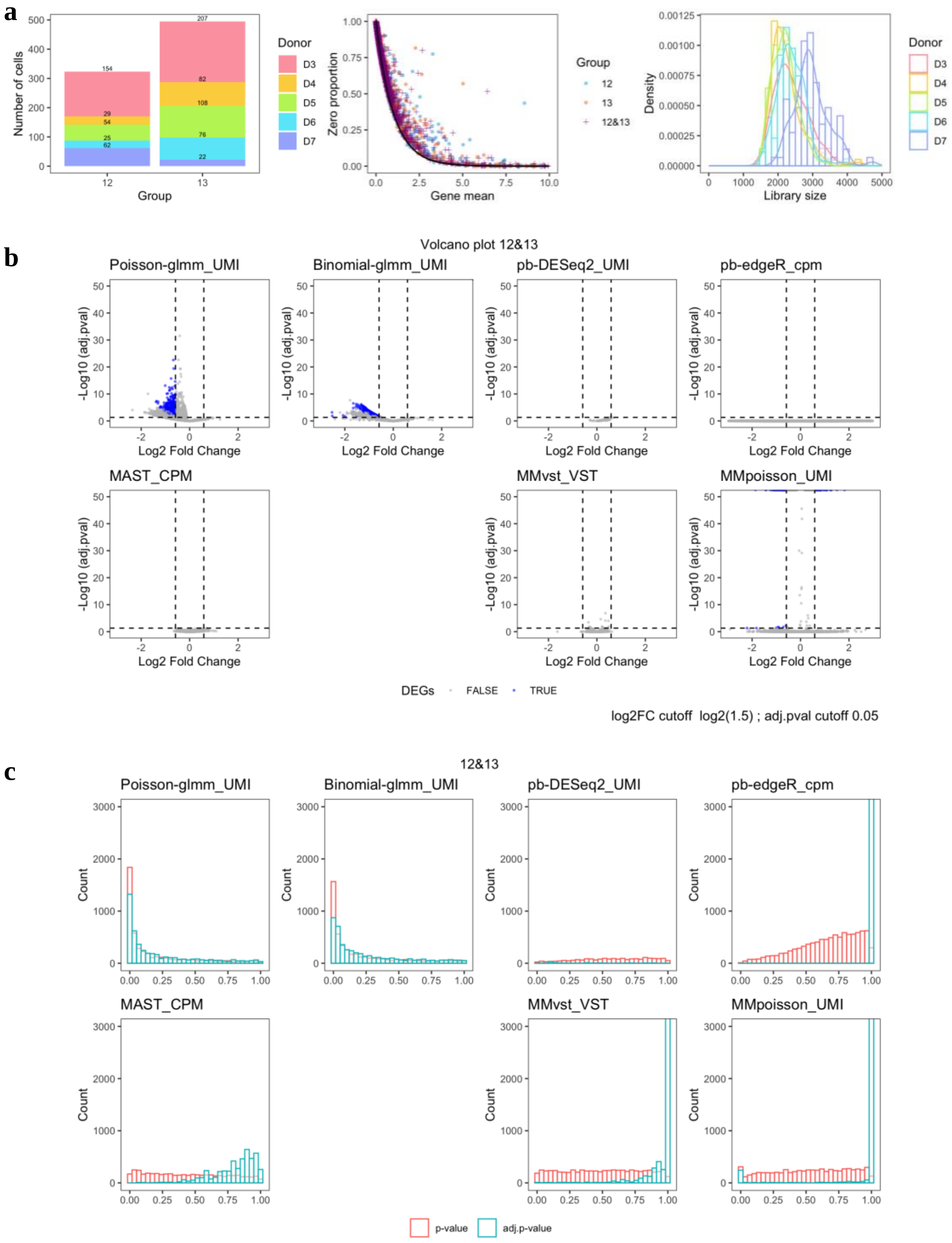
a
Number of cells
Group
Donor
D3
D4
D5
D6
D7
Zero proportion
Gene mean
12
13
12&13
Density
Library size
b
Volcano plot 12&13
Poisson-glmm_UMI
Binomial-glmm_UMI
pb-DESeq2_UMI
pb-edgeR_cpm
MAST_CPM
MMvst_VST
MMpoisson_UMI
-Log10 (adj.pval)
Log2 Fold Change
DEGs FALSE TRUE
log2FC cutoff log2(1.5) ; adj.pval cutoff 0.05
c
12&13
Count
p-value
adj.p-value

d

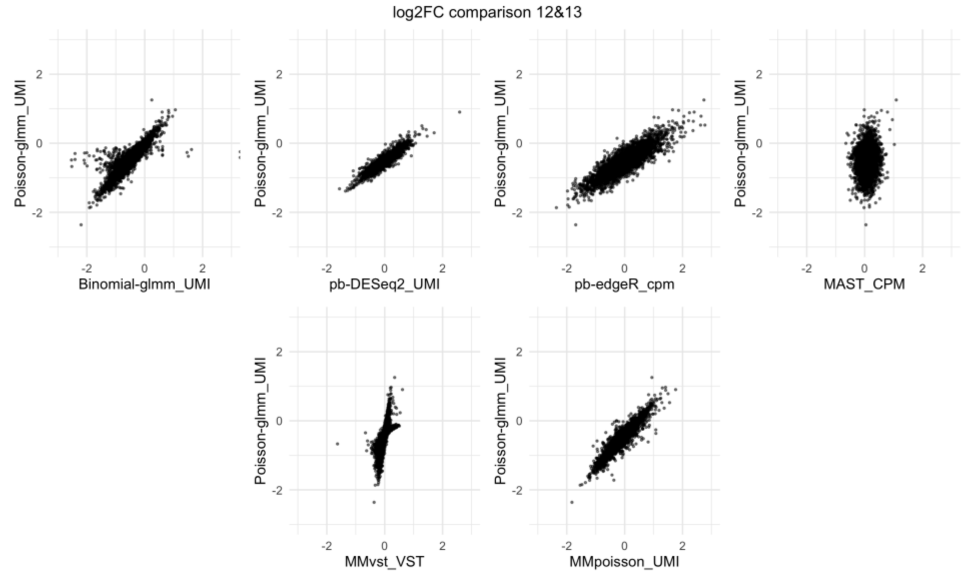


e

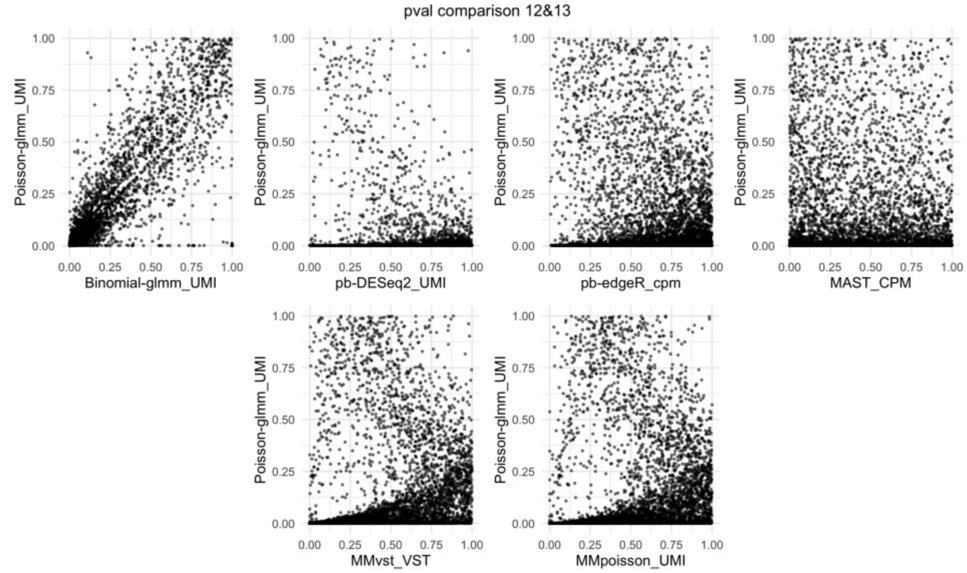


**Figure S3:** Additional diagnostic plots for group 12 and 13. **a)** Left: Donor composition in each group. Middle: Zero proportion plot for each group and combined. Right: Density plot of library size grouped by donors. **b)** Volcano plots for each method. Wilcox method is not applicable in this pair because the filtering procedure in Seurat excludes all genes. The signs of log2 fold change are adjusted such that positive signs represent higher expressions in group 13. **c)** Histogram of p-value and adjusted p-value for each method. **d)** Pairwise comparisons of log2 fold changes from other methods against LEMUR Poisson-glmm. **e)** Pairwise comparisons of p-values from other methods against LEMUR Poisson-glmm.

**Figure S4**

**a**

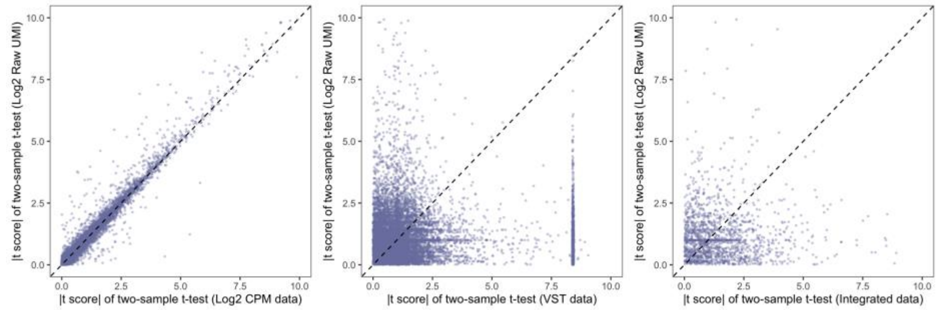


**b**

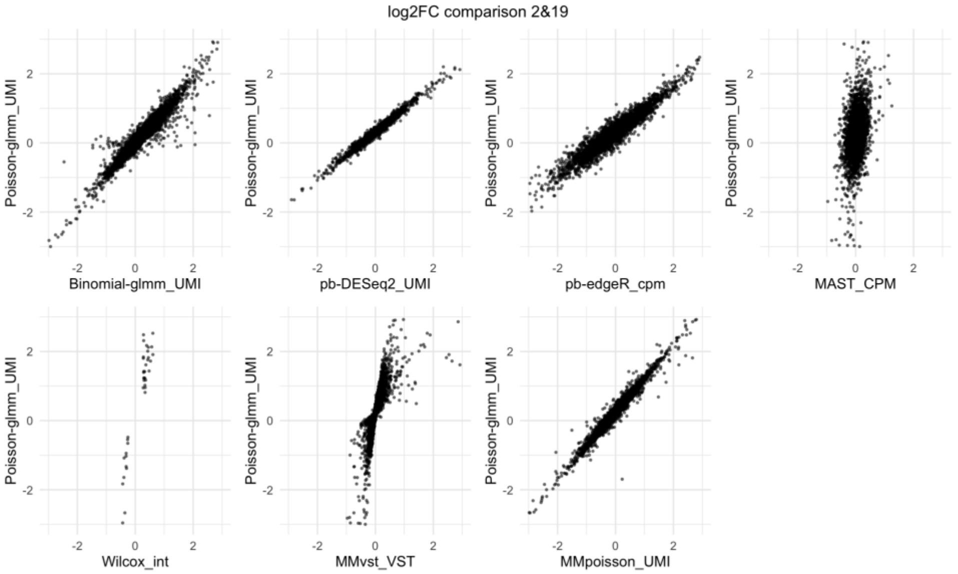


**c**

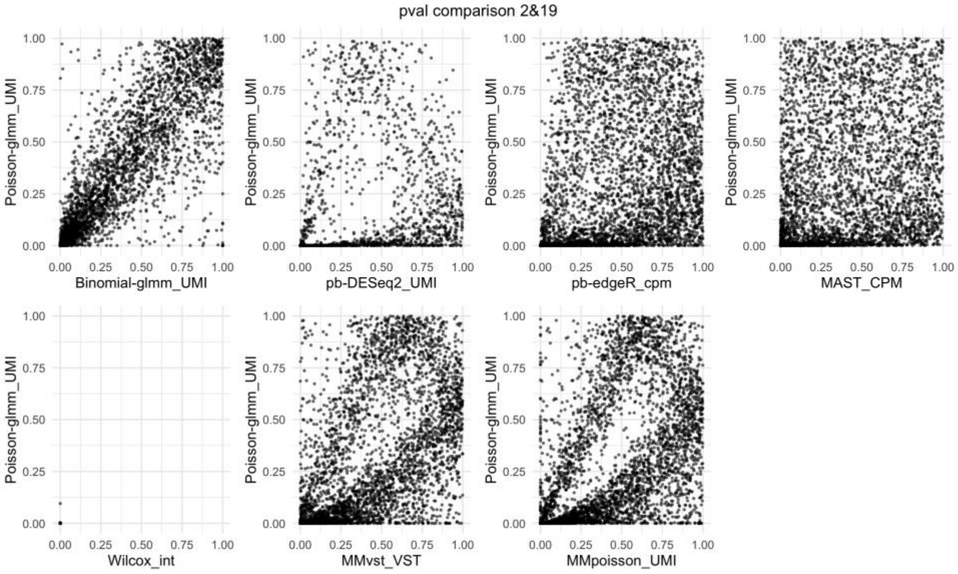

**d**

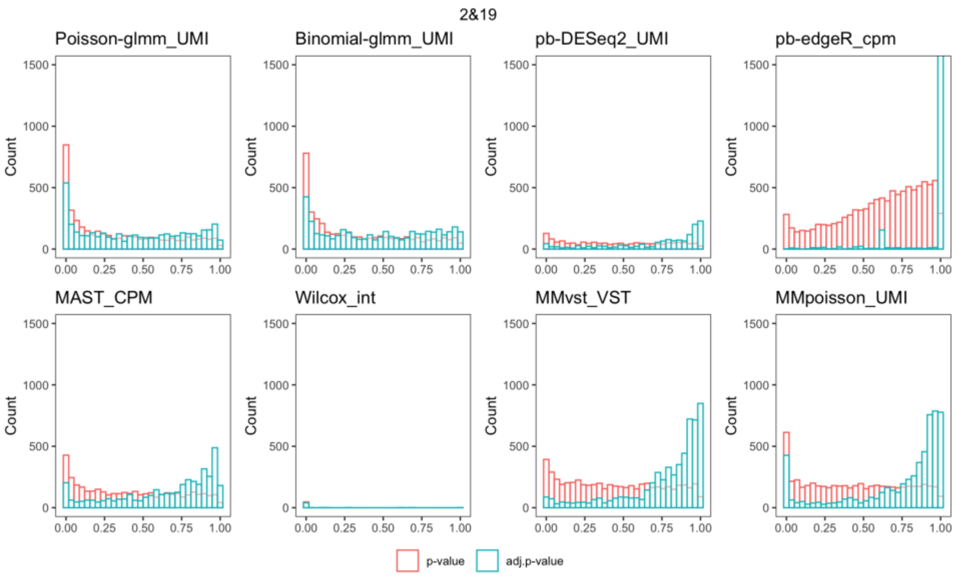
2&19
Poisson-glmm_UMI
Binomial-glmm_UMI
pb-DESeq2_UMI
pb-edgeR_cpm
MAST_CPM
Wilcox_int
MMvst_VST
MMpoisson_UMI
Count
p-value
adj.p-value

**e**

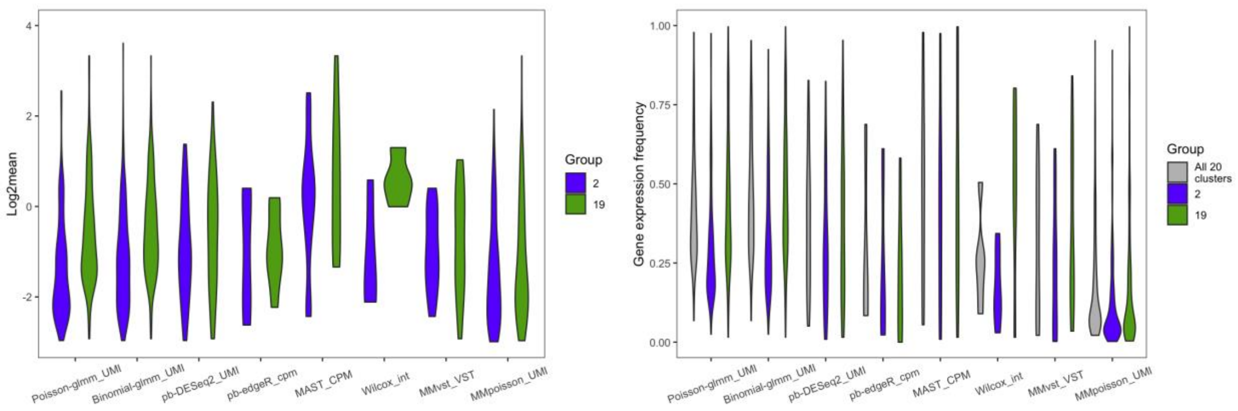
Log2mean
Group
2
19
Gene expression frequency
All 20 clusters
Poisson-glmm_UMI
Binomial-glmm_UMI
pb-DESeq2_UMI
pb-edgeR_cpm
MAST_CPM
Wilcox_int
MMvst_VST
MMpoisson_UMI

**f**

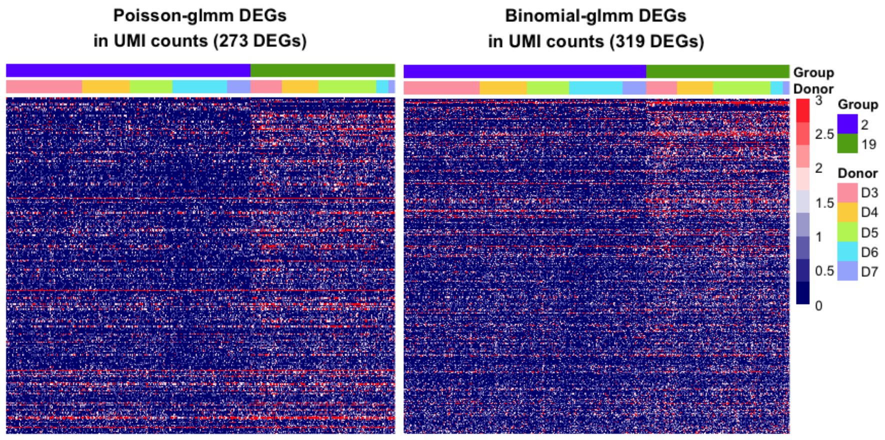
Poisson-glmm DEGs
in UMI counts (273 DEGs)
Binomial-glmm DEGs
in UMI counts (319 DEGs)
Group
Donor
2
19
D3
D4
D5
D6
D7

g

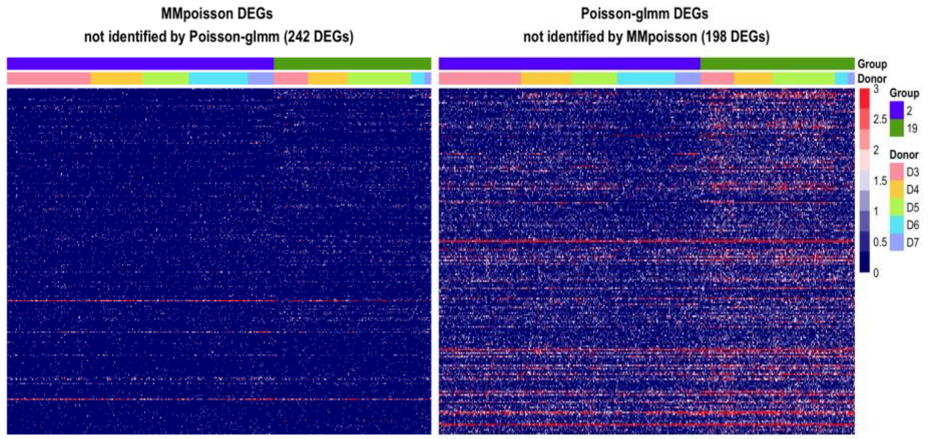


**Figure S4:** Additional diagnostic plots for group 2 and 19. **a)** Comparisons of t scores of mean difference test for raw UMI counts vs. other transformed data. **b)** Pairwise comparisons of log2 fold changes from other methods against LEMUR Poisson-glmm. **c)** Pairwise comparisons of p-values from other methods against LEMUR Poisson-glmm. **d)** Histogram of p-value and adjusted p-value for each method. **e)** Left: Violin plot of log2 gene mean for DEGs from different methods. Right: Comparisons of the gene expression frequency of the DEGs from different methods. **f)** Left: Heatmaps of DEGs from LEMUR Poisson-glmm. Right: Heatmaps of DEGs from LEMUR Binomial-glmm **g)** Left: Heatmaps of DEGs from MMpoisson but not identified by LEMUR Poisson-glmm. Right: Heatmaps of DEGs from LEMUR Poisson-glmm but not identified by MMpoisson.

**Figure S5**

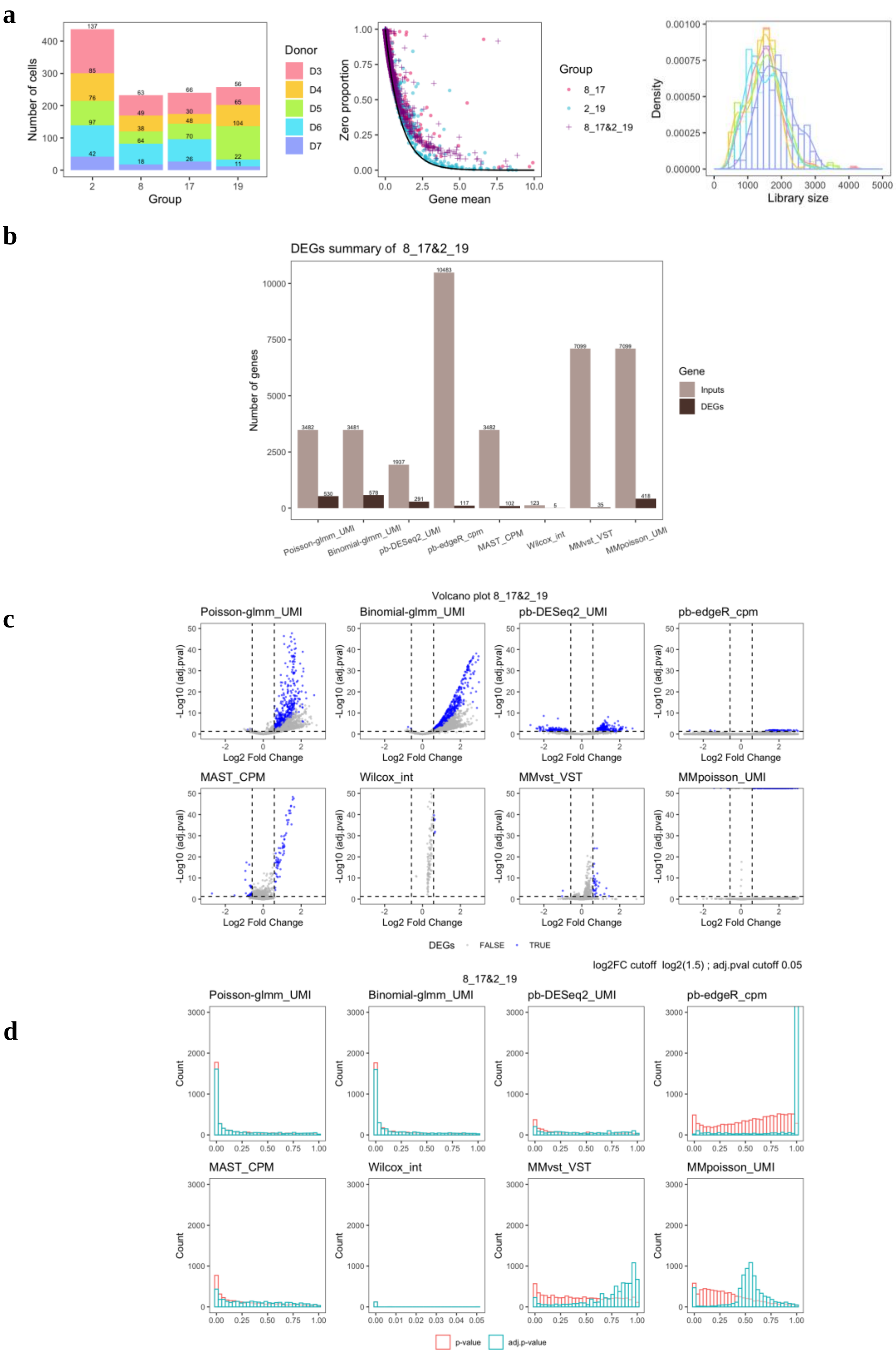

a
Number of cells
Group
Donor
D3
D4
D5
D6
D7
Zero proportion
Gene mean
8_17
2_19
8_17&2_19
Density
Library size
b
DEGs summary of 8_17&2_19
Number of genes
Gene
Inputs
DEGs
Poisson-glmm_UMI
Binomial-glmm_UMI
pb-DESeq2_UMI
pb-edgeR_cpm
MAST_CPM
Wilcox_int
MMvst_VST
MMpoisson_UMI
c
Volcano plot 8_17&2_19
-Log10 (adj.pval)
Log2 Fold Change
DEGs
FALSE
TRUE
log2FC cutoff log2(1.5) ; adj.pval cutoff 0.05
d
8_17&2_19
Count
p-value
adj.p-value

e

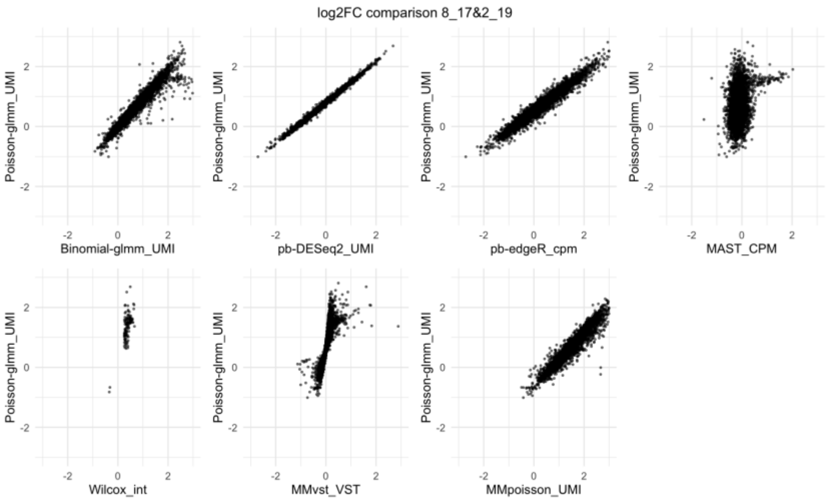


f

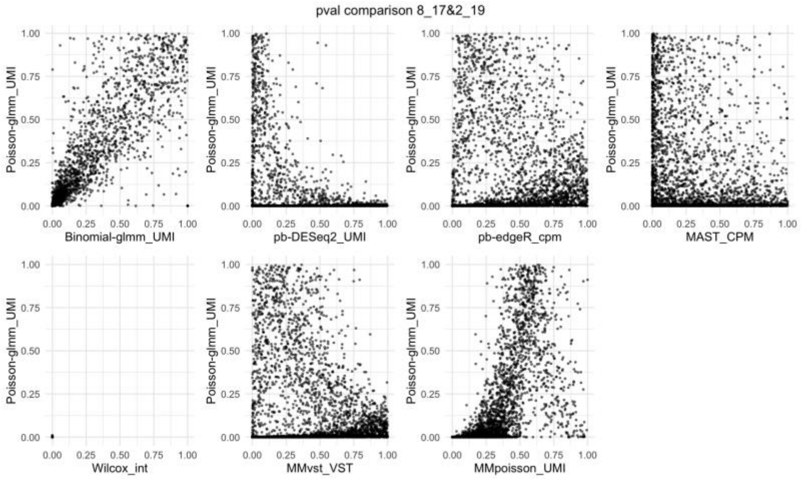


g

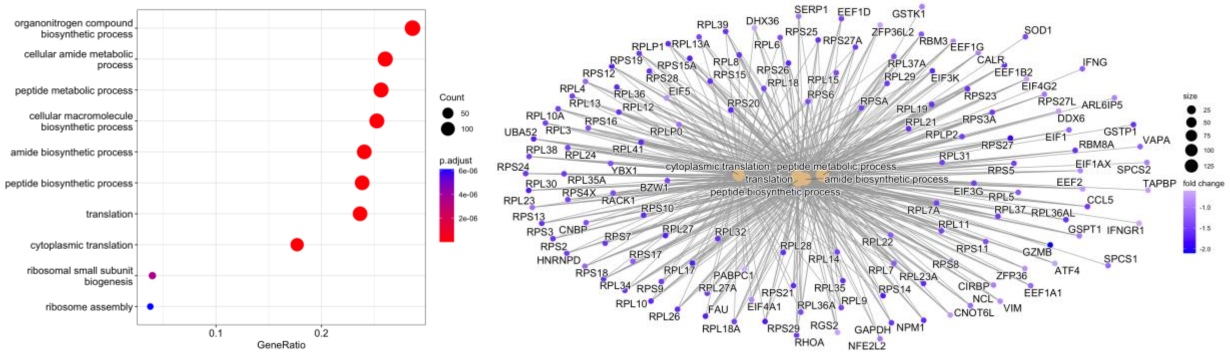


**Figure S5:** Additional diagnostic plots for group 8_17 and 2_19. **a)** Left: Donor composition in each group. Middle: Zero proportion plot for each group and combined. Right: Density plot of library size grouped by donors. **b)** Counts of input genes and DEGs in different DE methods. **c)** Volcano plots for each method. The signs of log2 fold change are adjusted such that positive signs represent higher expressions in group 2_19. **d)** Histogram of p-value and adjusted p-value for each method. **e)** Pairwise comparisons of log2 fold changes from other methods against LEMUR Poisson-glmm. **f)** Pairwise comparisons of p-values from other methods against LEMUR Poisson-glmm. **g)** GO analysis of the DEGs identified by Poisson-glmm.

**Figure S6**

**a**

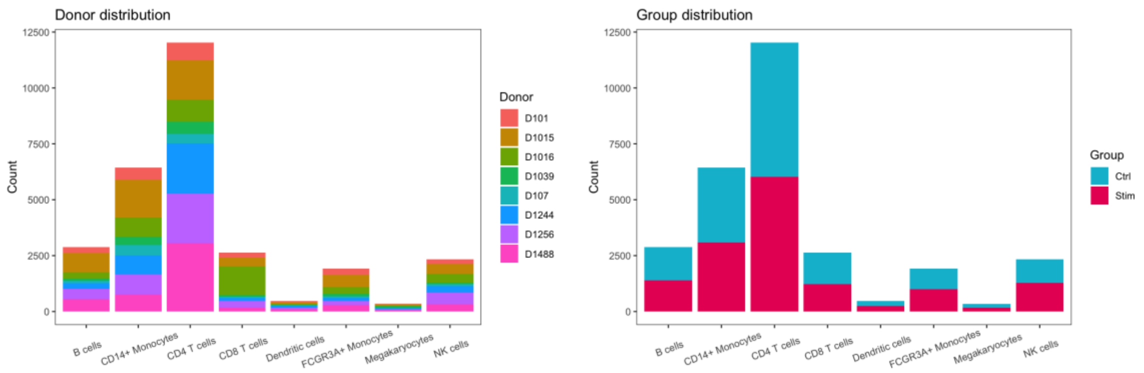


**b**

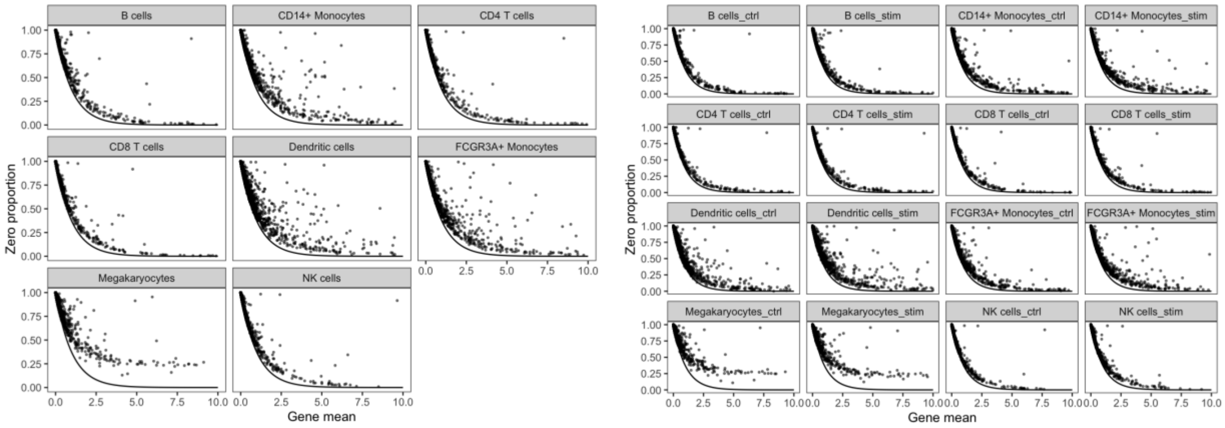


**Figure S6:** Additional data summary for case study 2. **a)** Left: Donor composition in each cell type. Right: Group composition in each cell type. **b)** Left: Zero proportion plots separated by cell types. Right: Zero proportion plots separated by cell types and group conditions.

**Figure S7**

**a**

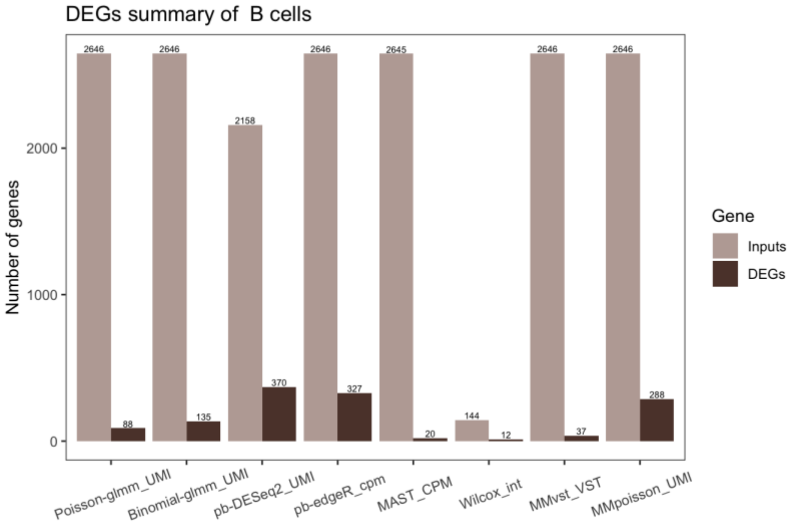


**b**

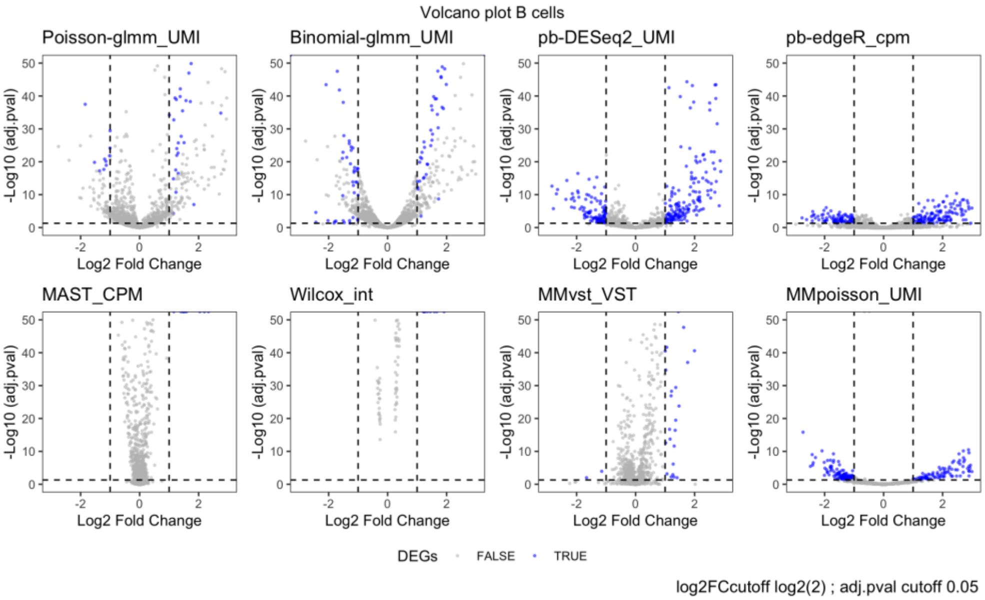


**Figure S7:** Additional diagnostic plots for B cells. **a)** Numbers of inputs and DEGs from different methods. Note that the input genes are all restricted to the input of Poisson-glmm **b)** Volcano plots for each method. The signs of log2 fold change are adjusted such that positive signs represent higher expressions in the stimulated group.

**Figure S8**

**a**

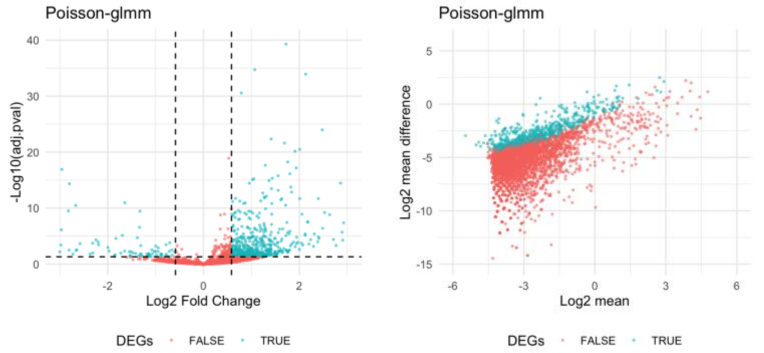


**b**

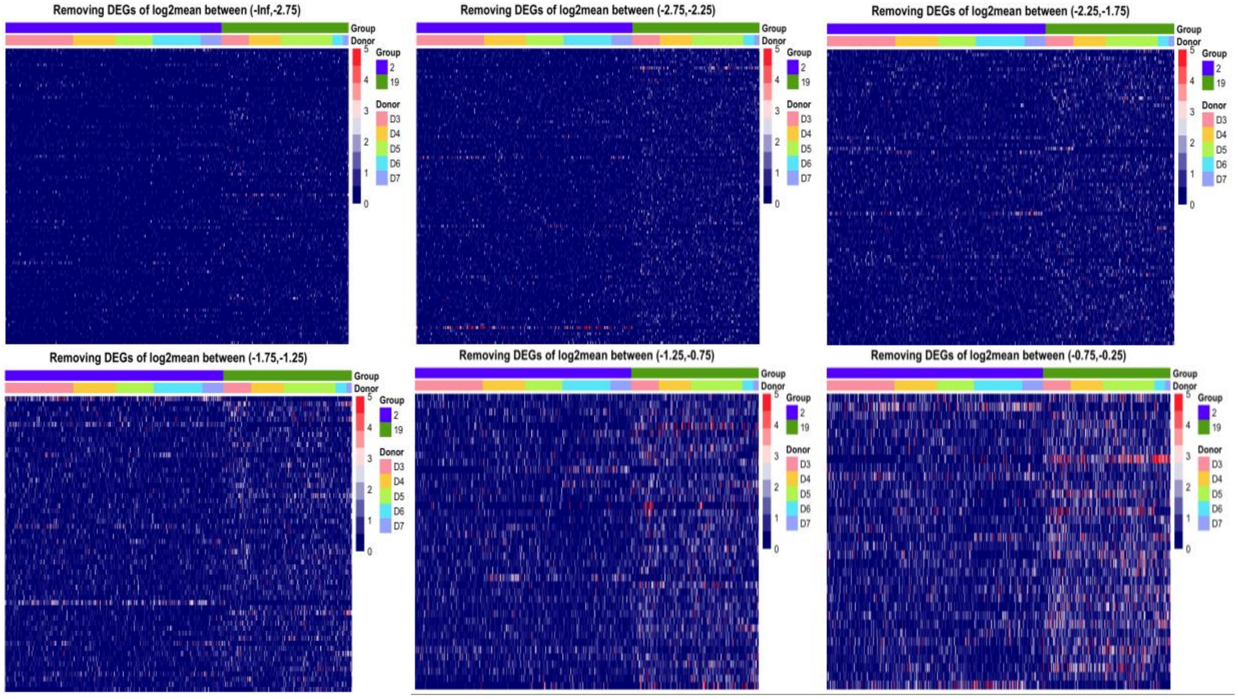


**c**

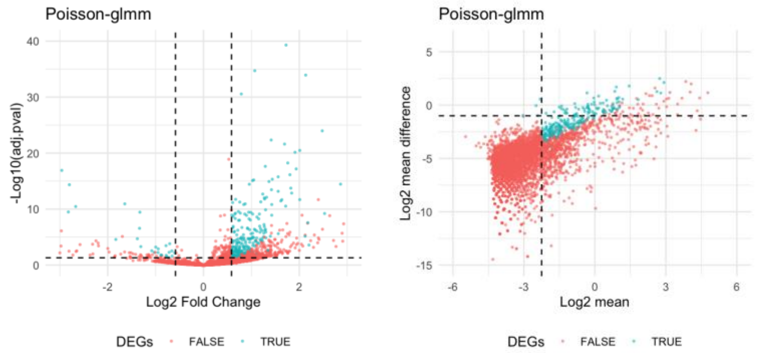


**Figure S8:** Diagnostic plots for determining DEGs. **a)** Left: Volcano plot for current criteria. Right: Gene mean vs. mean difference plot. **b)** Heatmaps illustrating the removed genes with small mean. **c)** Left: Volcano plot for new criteria. Right: Gene mean vs. mean difference plot. The DEGs selected by new criteria are annotated.

**Figure S9**

**a**

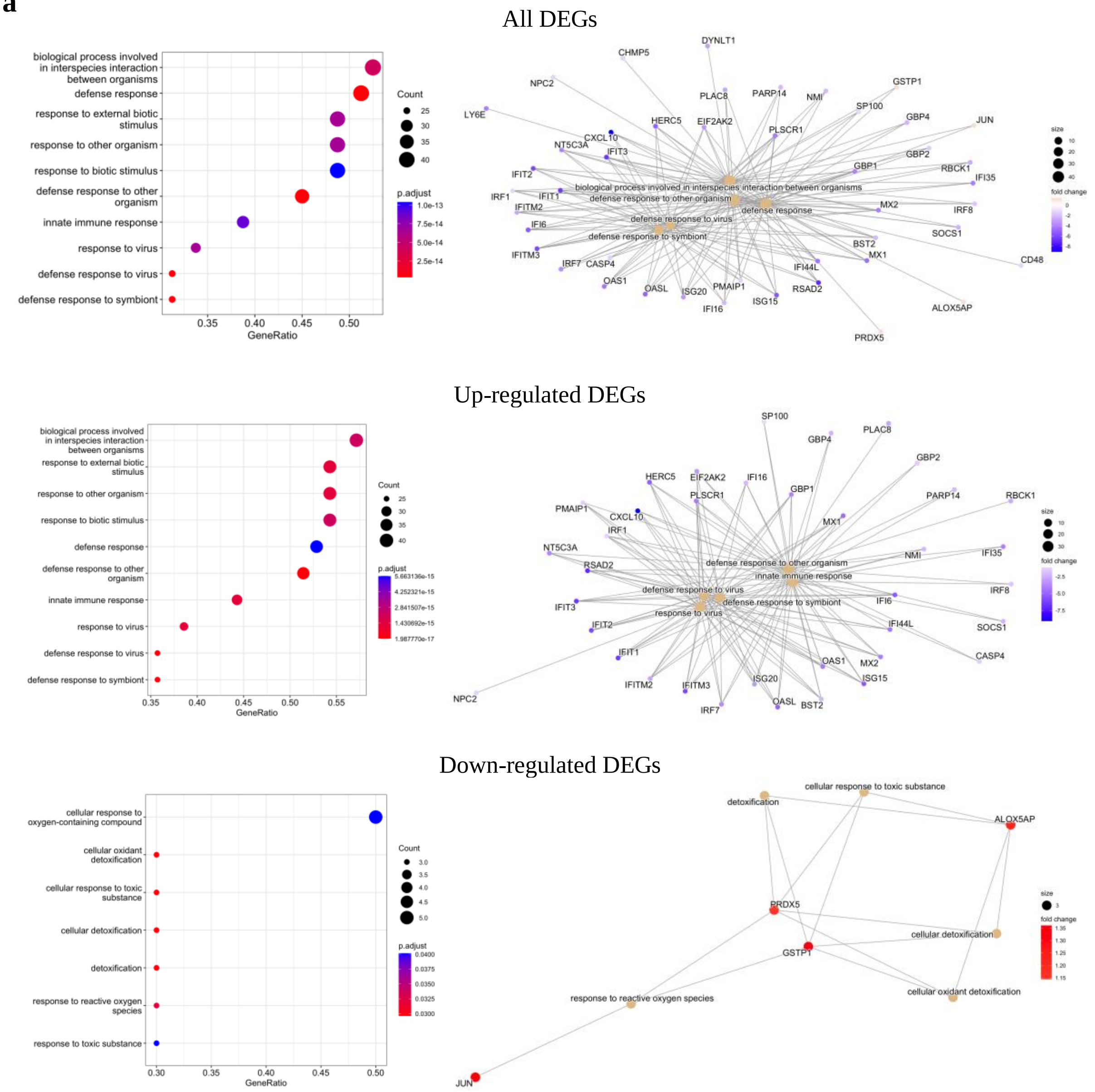
All DEGs
biological process involved in interspecies interaction between organisms
defense response
response to external biotic stimulus
response to other organism
response to biotic stimulus
defense response to other organism
innate immune response
response to virus
defense response to virus
defense response to symbiont
GeneRatio
Count
p.adjust
Up-regulated DEGs
Down-regulated DEGs
cellular response to oxygen-containing compound
cellular oxidant detoxification
cellular response to toxic substance
cellular detoxification
detoxification
response to reactive oxygen species
response to toxic substance
ALOX5AP
PRDX5
GSTP1
JUN
size
fold change

**b**

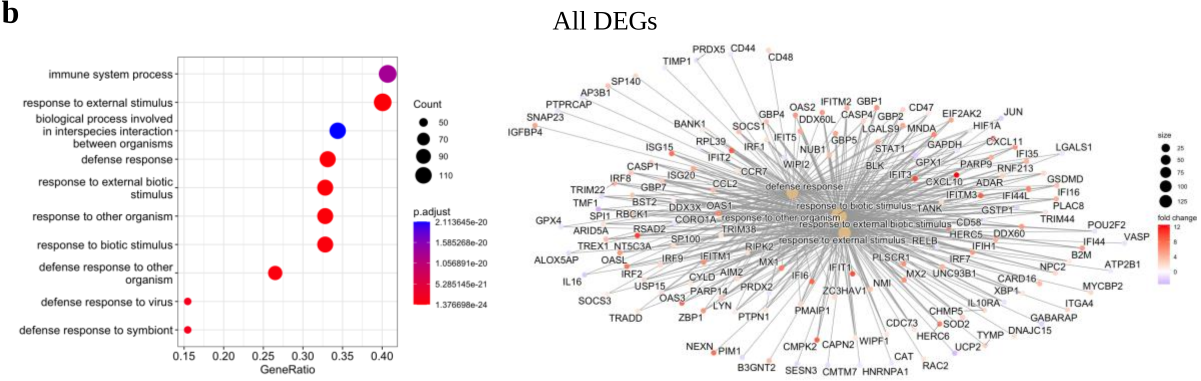
All DEGs
immune system process
response to external stimulus
biological process involved in interspecies interaction between organisms
defense response
response to external biotic stimulus
response to other organism
response to biotic stimulus
defense response to other organism
defense response to virus
defense response to symbiont
GeneRatio
Count
p.adjust
size
fold change

### Up-regulated DEGs

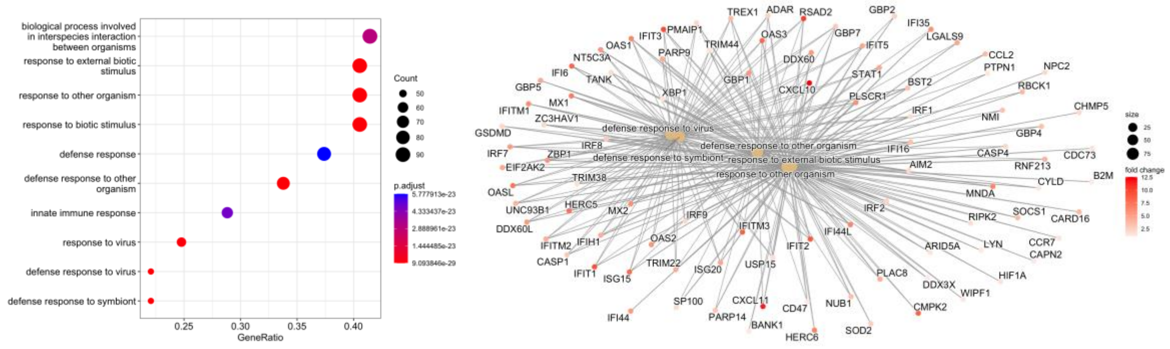


### Down-regulated DEGs

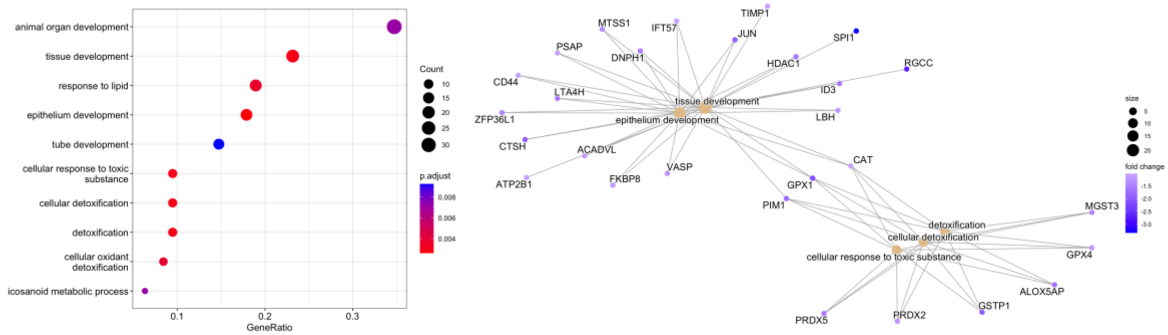


**c**

### All DEGs

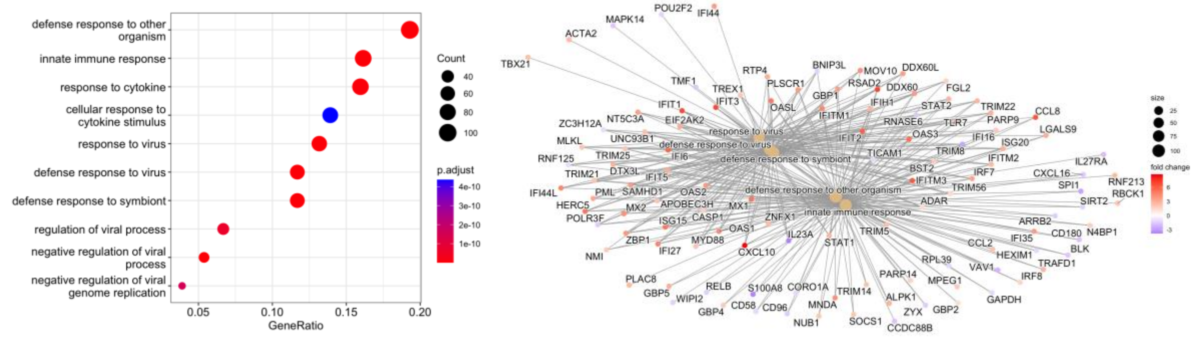


### Up-regulated DEGs

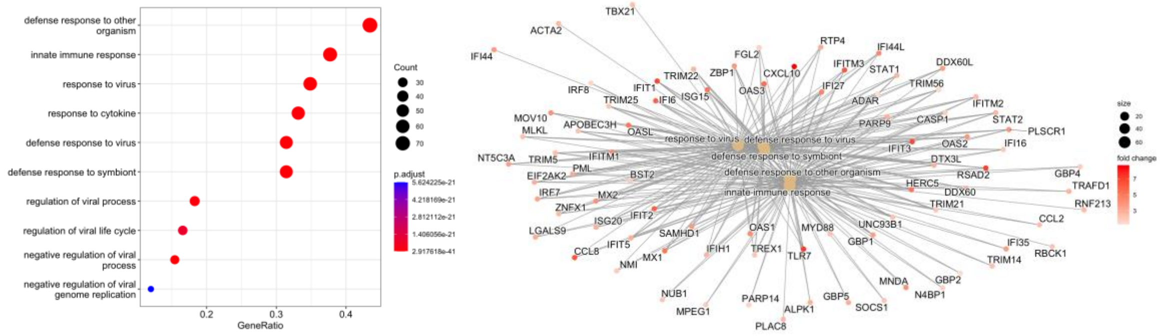

Down-regulated DEGs

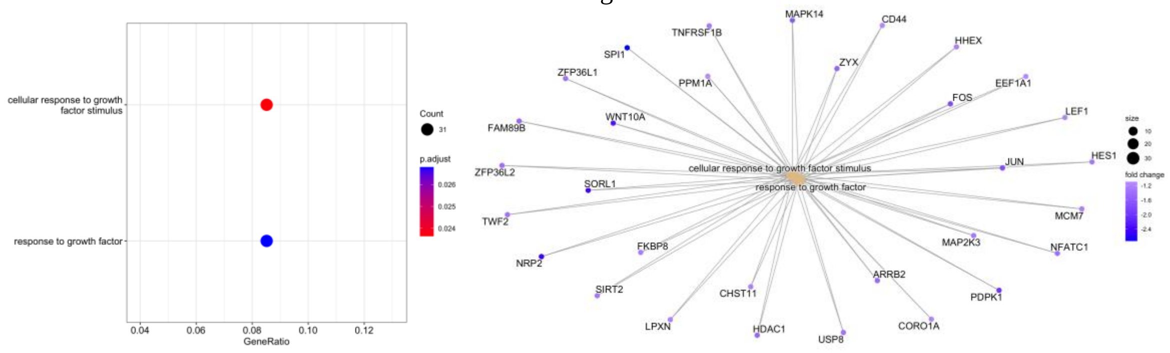


**Figure S9:** GO analysis of B cells by different DE methods. **a)** Poisson-glmm method. **b)** pb-DESeq2 method. **c)** MMpoisson method.

**Figure S10**

**a**

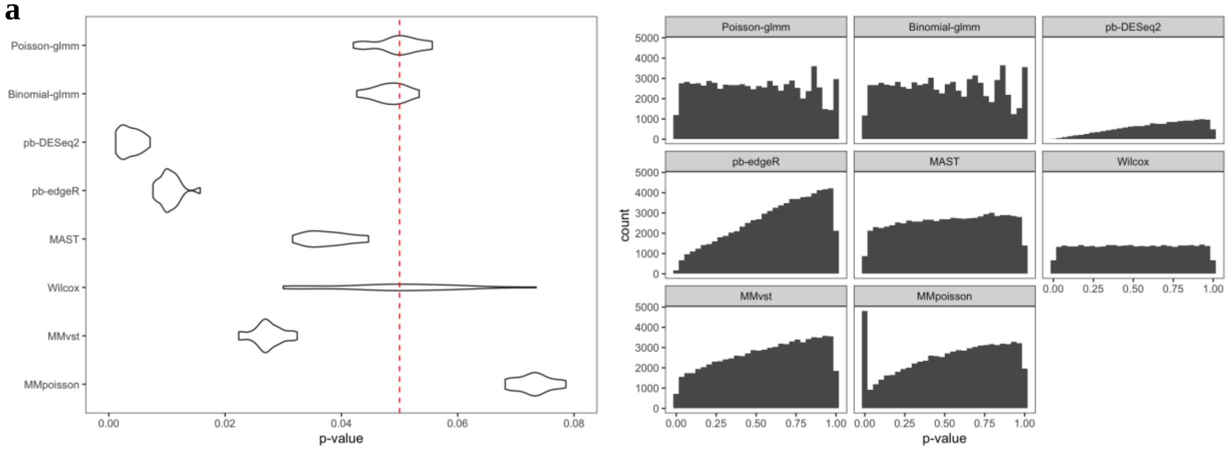


**b**

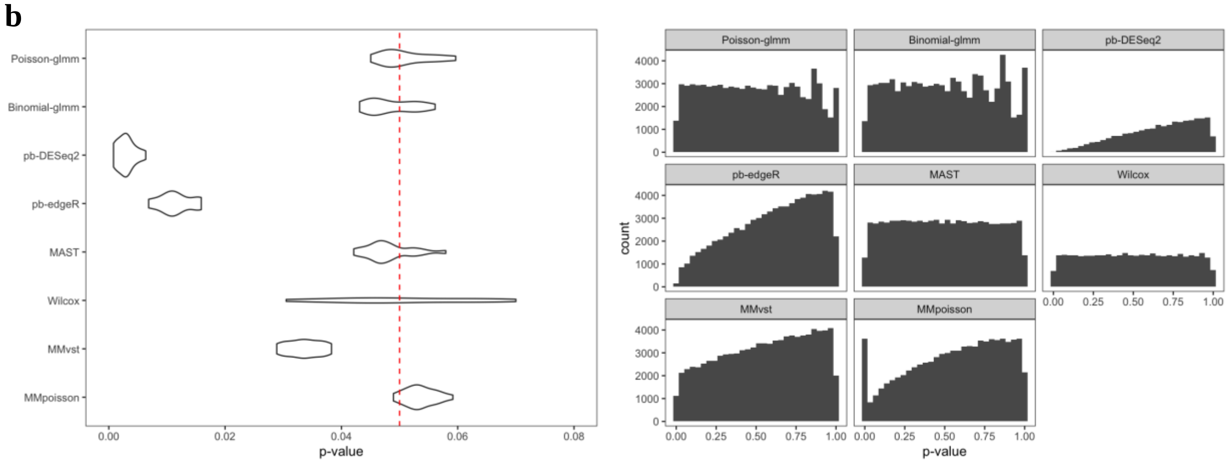


**c**

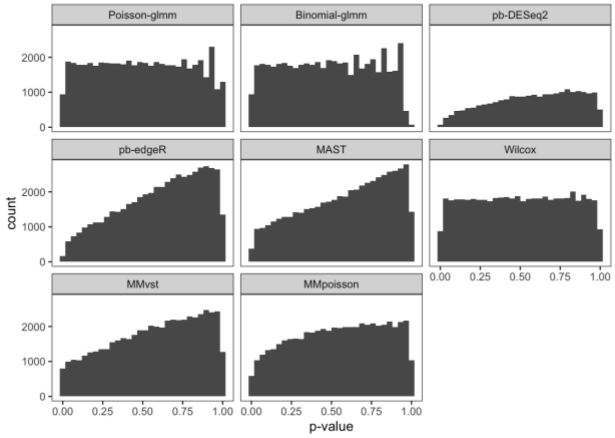


**Figure S10:** Permutation analysis under null setting on a dataset. **a)** Group 2 of case study 1. Left: Violin plots depicting the proportion of p-values below 0.05 for each method. Right: Histogram of p-values. **b)** Group 13 of case study 1. Left: Violin plots depicting the proportion of p-values below 0.05 for each method. Right: Histogram of p-values. **c)** Histogram of p-values under null setting on B cells of case study 2.
